# Competing 5’ splice sites of SRC1/HEH1 are marked for alternative splicing by U5 and U6 snRNAs during the B to Bact transition of the spliceosome

**DOI:** 10.1101/2025.06.06.658024

**Authors:** Ankita Katoch Banyal, Poulami Choudhuri, Balashankar R Pillai, Amjadudheen Varikkapulakkal, Shravan Kumar Mishra

**Affiliations:** Department of Biological Sciences, Indian Institute of Science Education and Research (IISER) Mohali, India; Institute of Molecular Biotechnology of the Austrian Academy of Sciences (IMBA), Vienna BioCenter (VBC), Dr. Bohr-Gasse 3,1030 Vienna, Austria

**Keywords:** Alternative 5’ splice site, Prp8, RES complex, Ecm2, U5 snRNA, U6 snRNA, B complex, Bact complex

## Abstract

Alternative splicing of precursor-messenger RNA (pre-mRNA) bearing introns with competing and overlapping 5’ splice sites (5’SS) produces more mRNAs. We investigated the mechanism of this form of alternative splicing by monitoring RNA and protein products of the yeast *SRC1/HEH1* gene. Its alternative splicing requires a sixteen-nucleotide pre-mRNA segment spanning the two 5’SS. The nucleotides are decoded by U5 and U6 small nuclear RNAs (snRNA), further supported by proteins of the spliceosomal B and Bact complexes. Specific variants of the pre-mRNA segment with recalibrated binding to U5 and U6 snRNAs allowed alternative splicing independently of the trans-acting proteins. Conversely, snRNA variants specified for modified 5’SS selection suppressed the splicing defects in the B complex mutants. Further data suggest that alternative splicing from *HEH1*-type introns containing overlapping and competing 5’SS relies on pre-mRNA-snRNA interactions during B to Bact transitions of the spliceosome, supported by proteins that stabilize low-fidelity, high-efficiency conformations.

## INTRODUCTION

Spliceosomes are large and dynamic ribonucleoprotein (RNP) complexes essential for constitutive and alternative splicing of precursor messenger RNAs (pre-mRNA). Each splicing cycle requires intricate rearrangements of five RNPs made of small nuclear RNAs (snRNAs) and their associated proteins (snRNPs). The snRNPs recognise and act through splicing signals of pre-mRNAs, including the donor 5’ splice site (5’SS) GU, the acceptor 3’ splice site (3’SS) AG, and the branch point (BP) adenosine. Two steps of the transesterification reactions, catalysed by divalent metal ions and U6 snRNA, remove introns and join exons to form functional mRNA^1, 2^. Alternative recognition of these signals can produce more mRNAs from a gene, thereby amplifying protein diversity from the limited gene pools in the organism^3, 4^.

Different classes of alternative splicing exist in eukaryotes. Key among them occurs through the selection of the alternative 5’SS^5^ and 3’SS^4^. Splicing through competing 5’SS is a prevalent mode of alternative splicing. A majority of pre-mRNAs in the budding yeast *Saccharomyces cerevisiae* use the highly conserved GUAUGU hexanucleotide donors^6^, but 5’SS sequences are highly variable in intron-rich eukaryotes, where alternative 5’SS selection is widespread^7^. Two major classes of 5’SS (NN/GURAG and AG/GUNNN), distinguished by the presence of ‘G’ at -1 and +5 positions, have been reported^8^. Alternative 5’SS are often found 4 nucleotides across the dominant 5’SS, and such occurrences in humans are estimated to be in the thousands^9, 10^. The canonical GU dinucleotide deviates to GC in 0.87% of the 5’SS^11^. The *HEH1/SRC1* gene *S. cerevisiae* possesses a similar alternative 5’SS.

Alternative splicing of *HEH1* pre-mRNA occurs through two non-canonical 5’SS hexanucleotides, /GCAAGU and /GUGAGU, arranged as a partially overlapping sequence: /GCAA/GUGAGU (bent lines indicate cleavage sites)^12^. The downstream 5’SS, /GUGAGU, is the dominant one and generates mRNA-encoding the longer Heh1L full-length protein, whereas splicing from the weaker upstream 5’SS /GCAAGU is tightly regulated and produces mRNA-encoding the shorter Heh1S protein (by gaining the tetranucleotide GCAA, *HEH1S* mRNA’s frame shifts at the exon-exon junction and acquires an in-frame stop codon in exon-2). The two proteins share a common N-terminus but differ in their C-termini, resulting in distinct topologies within the inner nuclear membrane. Heh1L is a double-pass membrane protein, with both its termini in the nucleoplasm, while Heh1S is a single-pass membrane protein with only the N terminus in the nucleoplasm^13^. Heh1 protein controls the expression of sub-telomeric genes^13^, anchors rDNA repeats^14^ and telomeric chromatin towards the nuclear periphery^13, 15^ and helps in chromatin silencing. Heh1S protein regulates Heh1L. The latter, if present alone, leads to the formation of aberrant nuclear pore complexes NPC^16^.

5’SS selection relies on early pre-mRNA recognition by trans-acting U1 snRNA and associated proteins^17–19^. The role of specific RBPs (RNA-binding proteins), SR (serine/arginine-rich) proteins, and hnRNPs (heterogeneous nuclear ribonucleoproteins) has also been reported in the context of 5’SS selection^5, 20–25^. These proteins act as splicing enhancers and silencers, and regulate spliceosome assembly on pre-mRNA targets in association with U1 snRNP. Later spliceosomal assemblies in 5’SS selection in humans have been recently reported^26^.

The *HEH1/SRC1* gene is an interesting tool for studying RNA and protein factors involved in alternative splicing through competing and overlapping 5’SS. Its alternative splicing is controlled by the ubiquitin-like protein Hub1 (UBL5 in humans) through non-covalent associations with the U1 and U2 snRNPs, bridging RNA helicase Prp5^27^ and the U4/U6.U5 tri-snRNP protein Snu66^28^. Thus, Hub1 acts early in spliceosome assembly by activating the RNA helicase, as well as later with the tri-snRNP component^29^. The latter activity suggests a possible role of the spliceosome core in *HEH1* 5’SS selection. Interestingly, we find that *HEH1* alternative splicing required optimal pairing of U5 and U6 snRNAs across a sixteen-nucleotide segment at the exon1-intron boundary of the pre-mRNA encompassing the competing and overlapping 5’SS. The 5’SS selection by the U5 and U6 snRNAs was supported by specific proteins of the spliceosome core, including Prp8.

## RESULTS

### Identification of trans-acting factors for *HEH1* alternative splicing

In the competing and overlapping 5’SS: /GCAA/GUGAGU of *SRC1/HEH1* pre-mRNA, the usage of the weaker upstream 5’SS: /GCAAGU was reported to require the ubiquitin-like protein Hub1 and its non-covalent binding partner Snu66 in the tri-snRNP complex. The dominant 5’SS: /GUGAGU was selected independently of Hub1^28^. However, the mechanism of 5’SS selection is not understood. We reasoned that by finding additional factors of this process, alternative 5’SS selecting spliceosomal complexes may be identified. Thus, we searched for *HEH1* alternative splicing factors in *Saccharomyces cerevisiae* through the following approaches.

i. *HEH1–LACZ alternative splicing reporters*: Along the line of *RP51-LACZ* splicing reporter^30^ we prepared *HEH1* alternative splicing reporters *HEH1S*–*LACZ* and *HEH1L*– *LACZ* (Figure S1a) by placing *HEH1* gene segment containing parts of both exons flanking the intron upstream of the β galactosidase-encoding *LACZ* gene such that yeast cells would produce in-frame translatable reporter only after precise 5’SS selections and excision of the intron. Reporter activities were monitored in cells growing on solid and liquid media. Besides the Hub1 mutant, reduced activity from the alternative 5’SS was also observed in the deletion mutants of the RES^31^ (Retention and Splicing complex) complex subunits Snu17, Bud13, Pml1, and Urn1. The deletion of the NTC (Nineteen complex) factor Ecm2^32^ strain also showed similar defects in 5’SS selection (Figure S1b and c). By contrast, the usage of the dominant 5’SS remained unaltered in these mutants, suggesting a role of these proteins in the selection of the weaker 5’SS.
ii. *Heh1 protein isoforms*: The relative abundance of the two Heh1 protein isoforms, Heh1S and Heh1L, was monitored by western blot assays. *HEH1* gene was either ectopically expressed under a heterologous promoter from a plasmid, or chromosomally epitope-tagged at the N-terminus, in deletion and point mutants of splicing factors. Complementing the findings from the reporter assays, Heh1S protein was strongly diminished in the knockout mutants of Snu17, Bud13, Pml1, and Urn1, in addition to Hub1 (Figure S1d). The loss of potential RNA-binding residues of Hub1, as deduced from cryo-EM studies of spliceosomes^33, 34^, showed a similar lack of Heh1S protein (Figure S1h). As expected, the previously reported D22A and H63L mutants of Hub1 were defective in *HEH1* alternative splicing. Hub1-binding deficient Snu66-HIND mutants also showed a lack of Heh1S (Figure S1j). A set of distinct Prp8 alleles was screened for alternative splicing defects by expressing *HEH1* gene from a plasmid with an N-terminal epitope tag. Interestingly, Heh1S protein was diminished specifically in the *prp8-101* (E1960K) allele^35^isoforms in mutants identified above are placed in (Figure 1a). While Heh1S protein was strongly diminished in all the mutants than the wt strain, Heh1L isoform was higher in the mutants (Figure 1a), possibly because the lack of competition from the *HEH1S* 5’SS site led to preferential usage of the dominant *HEH1L* 5’SS
iii. *HEH1 mRNA isoforms*: *HEH1* transcripts were monitored by reverse-transcription (RT) PCR assays with primers binding in *HEH1* exons. More retention of intron was observed in Δ*hub1* and Δ*snu66* mutants compared to Δ*ecm2, prp8-101*, and the RES factor mutants (Figure 1b). The data indicated normal constitutive splicing and enhanced usage of the dominant 5’SS in the absence the competing one in the mutants. We verified these observations by purifying and sequencing *HEH1* cDNAs from the mutants. Since *HEH1S* and *HEH1L* mRNAs differ in GCAA nucleotides at their exon junctions (*HEHL* gains them due to the usage of downstream 5’SS), the electropherogram shows two mixed peaks after the common exon. Areas under the peaks were integrated to estimate the relative abundance of the two mRNAs. Consistent with the above findings from reporter assays and protein analysis, the mutants showed diminished levels of *HEH1S* mRNA, confirming the 5’SS selection defects at the RNA level (Figure 1c). The electropherograms further showed that *HEH1L* mRNA level was more in the mutants compared to the wt strain (Figure 1d), consistent with higher protein levels seen in Figure 1a. Thus, spliceosomes used the dominant *HEH1L* 5’SS more efficiently in the absence of competition from *HEH1S* 5’SS.
iv. *Probing hub1 proximity to splicing factors*: In cells deleted of Hub1, Snu66-binding-deficient *hub1(D22A)* mutant in free form does not support the usage of *HEH1* alternative 5’SS, but its linear fusion to Snu66, Prp38 and Prp8 restored alternative splicing^28^. We used this strategy to find Hub1’s proximity to splicing factors essential for cell viability, which could not be analysed in the above assays. Interestingly, *hub1(D22A)* fused to Prp3, Prp6, and Prp31 visibly restored *HEH1* alternative splicing (Figure 1e). Notably, proximities to only specific proteins of the spliceosome B^36^ and Bact complexes^37^ (Snu66, Prp38, Prp8, Prp3, Prp6, and Prp31), but not to other proteins of the B complex or the proteins of U1 and U2 snRNPs, restored alternative splicing (Figure 1e).

**Figure 1:**
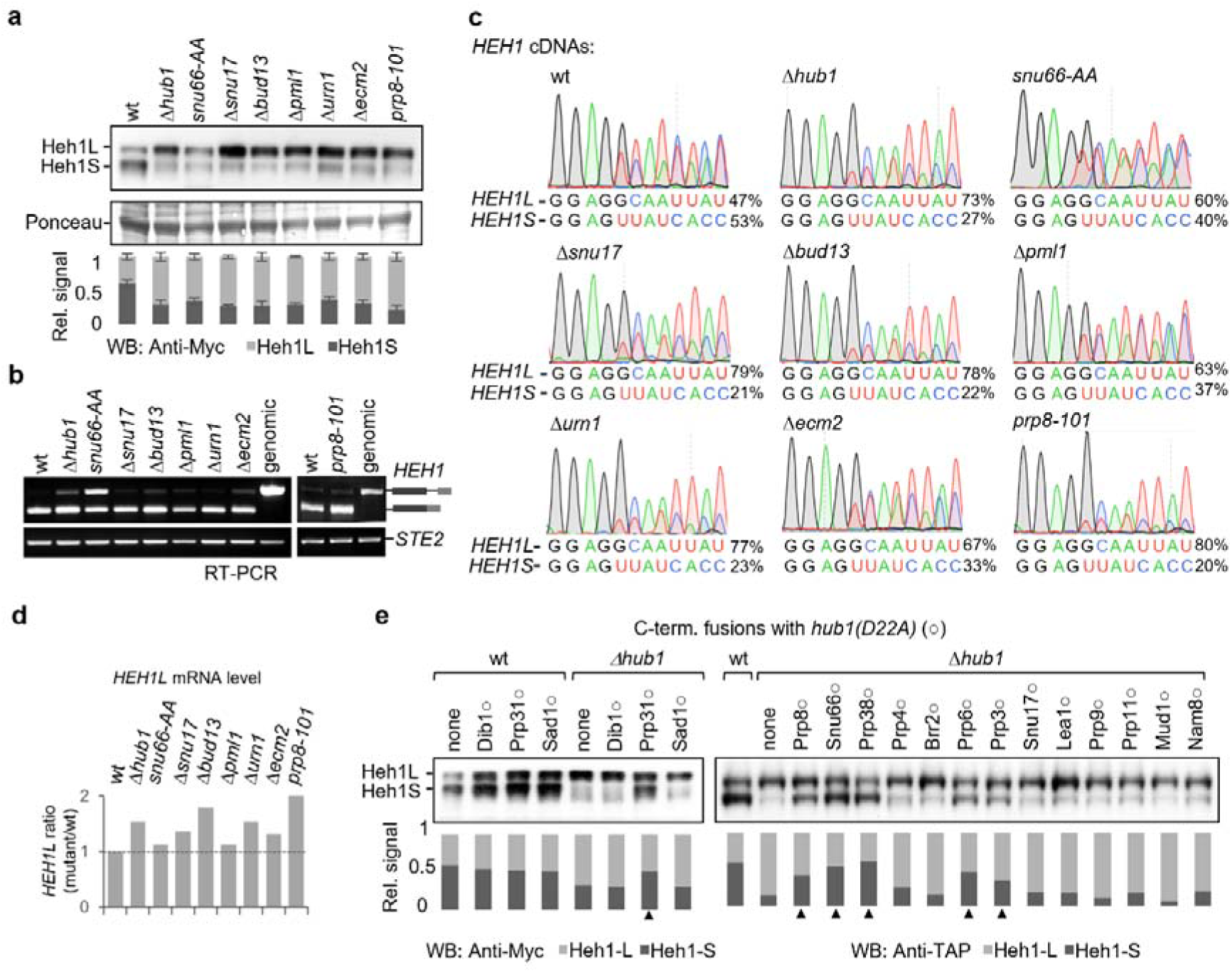
*HEH1* alternative splicing needs B and Bact proteins. **(a)** The abundance of Heh1 L and S protein isoforms in the mutants of indicated spliceosomal proteins was monitored by western blots. Data shown as mean ± s.d. (n=3 independent replicates). **(b)** RT-PCR assays to monitor of *HEH1* pre-mRNA splicing in deletions and mutant strains of indicated splicing factors. *hub1*, *snu17*, *ecm2*, and *urn1* mutant strains are respective gene knockouts. *snu66-AA* mutant refers to R16A and R47A variant of Snu66, its Hub1-binding-deficient HIND mutant. *prp8-101* is the E1960K allele of Prp8. The mutants were identified from different targeted screens shown in Figure S1a-f, h-j, and k-l. **(c)**Spliced cDNA bands were purified from mutants in (b) and sequenced to estimate the extent of *HEH1* alternative splicing from the area under peaks of isoform-specific nucleotides. The values for UU<u>A</u>UCACC indicate the relative abundance of *HEH1S*, and GC<u>A</u>AUUAU shows the relative abundance of *HEH1L*. Peak for the underlined A, being common to both isoforms, were omitted from the quantitation. The values on the right of the peaks indicate relative % abundance of respective mRNA isoforms. **(d)** Relative abundance of *HEH1L* mRNA in different strains was estimated from cDNA electropherogram (shown in Figure 1c) **(e)** Splicing factors from distinct spliceosomal complexes were probed by linear chromosomal fusions of a Snu66-binding-deficient *hub1(D22A)* mutant, indicated with **○**, at the C-termini of respective splicing factors. Rescue of alternative splicing defects (Heh1S protein) in the *Δhub1* background following *hub1(D22A)* fusion was monitored in western blots. Upward triangle (▲) marks *Δhub1* strain harboring splicing factor-*hub1(D22A)* fusions, where Heh1S protein were restored. Note that the otherwise free *hub1(D22A)* is deficient in *HEH1* alternative splicing^28^ (see Figure S1h).

The data discussed above in (i-iv) confirmed the role of specific proteins of the spliceosome core in *HEH1* alternative splicing. Interestingly, these factors did not lead to dominant usage of the alternative 5’SS. Overexpression of none of these proteins forced 5’SS selection towards the weaker 5’SS or suppressed the splicing from the dominant 5’SS (Figure S1m-n). Even double chromosomal fusions of *hub1(D22A)* to both Snu66 and Prp38 did not enhance splicing from the weaker 5’SS or suppress the dominant 5’SS (Figure S1o). Thus, the trans-acting proteins play critical roles in 5’SS selection, but do not dominate over the cis elements in the pre-mRNA. These findings raised an interesting question: if not by proteins, how are the competing 5’SS decoded?

### U5 and U6 snRNAs decode exon-intron boundaries for alternative 5’SS selection

From alternative splicing reporters discussed above and known binding sites of U1, U5, and U6 snRNAs at exon-intron boundaries, we narrowed a 16-nucleotide pre-mRNA segment (1913-1928 of *HEH1* gene) critical for alternative 5’SS selection (Figure 2a). To study their importance, *HEH1* expression constructs containing changes at every nucleotide (except for the 5’ donors ‘G’) were made by site-directed mutagenesis. Proteins originating from the pre-mRNA variants were studied by western blots (Figure 2b), and mRNA isoforms by cDNA sequencing (Figure 2C). Changes resulting in stop codons could be studied only by cDNA sequencing (for example, G<u>C</u>AA to G<u>U</u>AA change gains an in-frame stop codon in *HEH1L* mRNA, explaining why a GC donor instead of the common GU is used in *HEH1*). Almost any change in this region produced either Heh1S or Heh1L protein (barring the *HEH1S*•{G(−1)A} variant 4). These results highlighted the importance of precisely selected nucleotides around competing 5’SS for alternative splicing.

**Figure 2:**
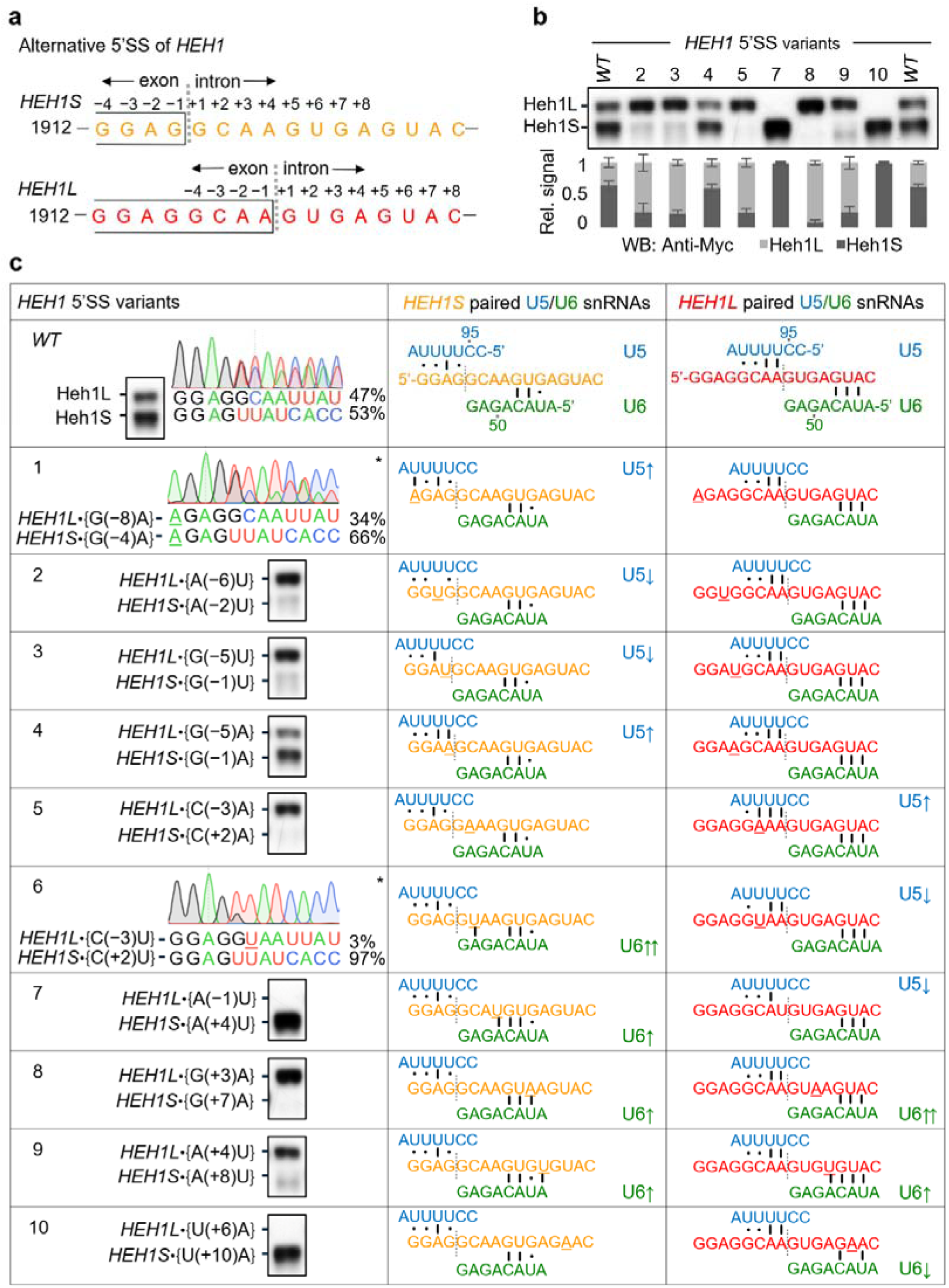
Sequence at the exon-intron boundary determines alternative 5’SS selection. **(a)** The 16 nucleotides segment at the exon/intron (1913-1926) boundary of *HEH1* was investigated for alternative splicing. The sequence was selected based on reporter assays described earlier, pairing of U5 snRNA loop1 and U6 snRNA ACAGA box. *HEH1S* (orange color) and *HEH1L* (red color) sequences represent isoforms of *HEH1* pre-mRNA. **(b)** *HEH1* was mutated sequentially from left to right and alternative selection was monitored by western blot. Data shown as mean ± s.d. (n=3 independent repeats). **(c)** Analysis of pre-mRNA variants paring with U5 snRNA (Blue) and U6 snRNA (Green) for both *HEH1S* and *HEH1L* was tabulated. The mutants were analyzed for presumed U5 and U6 snRNA pairings adapted from^37, 62^. Western blot lanes are taken from (b). The number in the left column shows variants ID. Resulting changes for *HEH1S* and *HEH1L* are shown in curly brackets {}. Filled dots (●) shows non-Watson-Crick pairing, standing lines (**ı**) represent Watson-Crick pairing, upward arrows (↑) mean increase in U5 / U6 pairing compared to WT *HEH1S* or *HEH1L*, downward arrows (↓) show decrease in U5 / U6 pairing compared to WT. and double upward arrows (↑↑) represent the yeast canonical 5’SS (GUAUGU or GUAAGU. The variants leading to premature stop codons, indicated by asterisk (*), could be analysed only by cDNA sequencing. The values on the right of the peaks indicate relative % abundance of respective mRNA isoforms.

The loss of alternative splicing of the pre-mRNA variants, however, did not correlate with U1 snRNA-5’SS interaction. Strikingly, we noticed a perfect correlation in the 5’SS choice and base-pairing strengths with U5 and U6 snRNAs. 5’SS that paired stronger to U5 or U6 snRNAs appeared to be preferentially used as the donor site and led to the loss of splicing from the second site. These observations underlined the role of the spliceosome core with U5 and U6 snRNAs in 5’SS selection. These results also suggested potential competition of the two 5’SS for binding to U5 and U6 snRNAs, and only optimal pairing strengths would allow selection of both 5’SS. Extending this idea, U5 and U6 snRNA variants with modified pairing strengths should change 5’SS selection. We tested this possibility by overexpressing U1, U5, and U6 snRNA variants with reinforced pairing to the weaker 5’SS, and monitored the alternative splicing pattern by RNA and protein assays (Figure 3b-c and Figure S3a-d).

**Figure 3:**
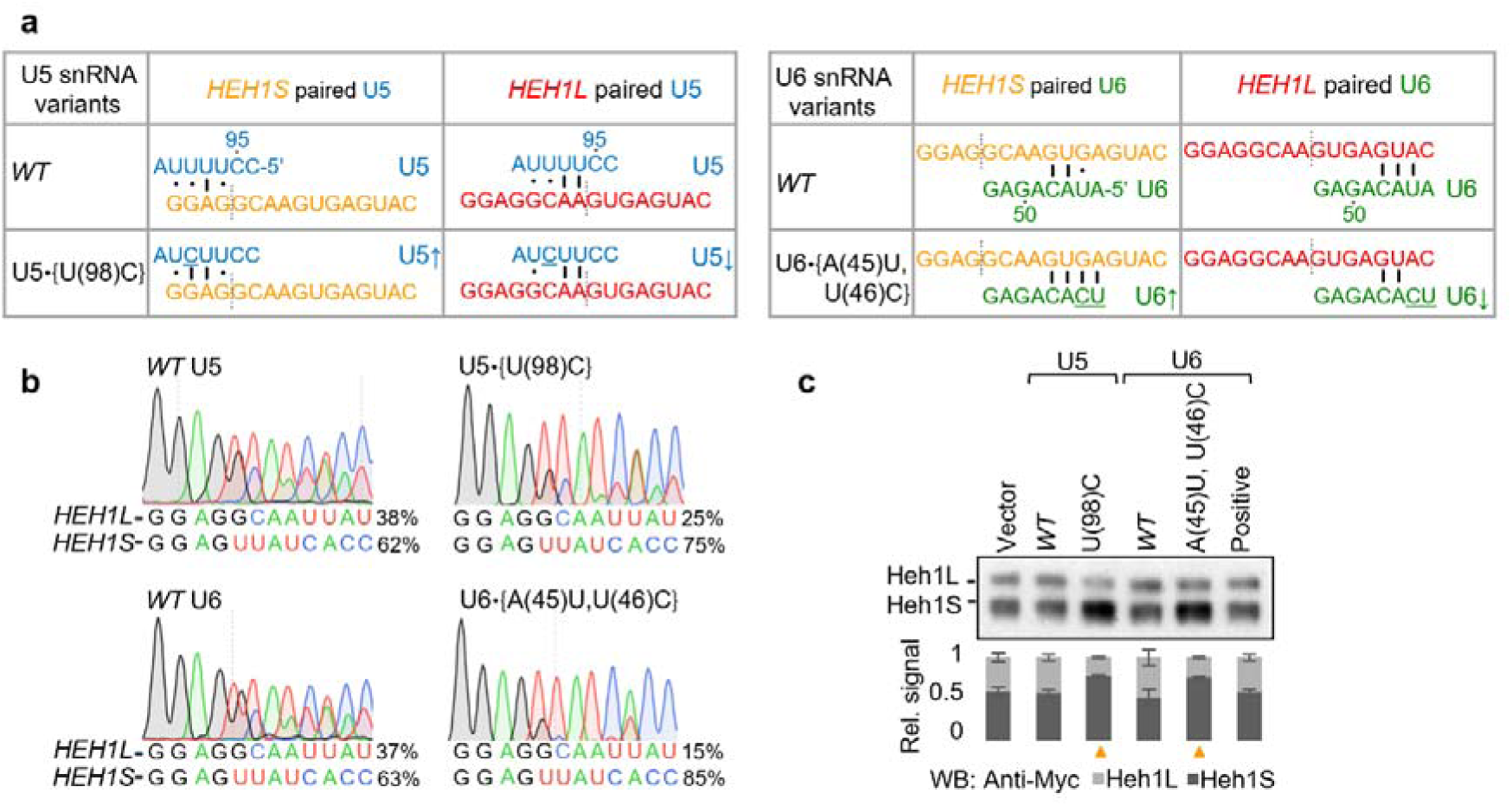
Role of U5/U6 pairing in *HEH1* alternative splicing. **(a)** The pairing of U5 snRNA and its variant U5•{U(98)C} and U6 snRNA and its variant U6•{A(45)U,U(46)C} to *HEH1* pre-mRNAs. The snRNA variants were identified (Figure S3a-b and S3c-d) by mutagenesis of U5 Loop1 and U6 ACAGA box. Up/down arrows (↑/↓) refer to increase/decrease in U5/U6 pairing to *HEH1* isoforms. **(b)** *HEH1* cDNAs from yeast strains overexpressing U5 and U6 snRNAs and their variants was sequenced to monitor alternative 5’SS selection. Area under the peak was integrated (similar to Figure 1) the find the relative abundance of the two mRNA isoforms.The values on the right of the peaks indicate relative % abundance of respective mRNA isoforms. **(c)** The expression of Heh1 protein isoforms upon overexpression of U5 and U6 snRNA variants monitored by western blots. Data shown as mean ± s.d. (n=3 independent replicates). Orange upward triangle (▴) indicates snRNA overexpression resulting in relatively higher amount of Heh1S protein.

Strengthened pairing of U5 and U6 snRNAs to a 5’SS favoured its selection. Stronger pairing of U(98)G variant of U5 snRNA and A(45)G variant of U6 snRNA to *HEH1L* 5’SS produced more Heh1L protein (Figure S3b and S3e). On the other hand, stronger pairing of U(98)C variant of U5 snRNA and U(46)C A(45)U variant of U6 snRNA variants to *HEH1S* 5’SS induced its preferential selection and produced more *HEH1S* mRNA and Heh1S protein (Figure 3b-c and S3b and S3d). Furthermore, pairing of 46^th^ and 45^th^ nucleotides of U6 snRNA, beyond the ACAGA box, to +7 and +8 nucleotides of the intron are crucial determinants for choosing a competing 5’SS (Figure 3b-c and Figure S3d). As discussed in Figure S1, overexpression of the protein factors did not induce a similar bias in 5’SS selection. Thus, optimal decoding of competing 5’SS by U5 and U6 snRNAs appears to be the primary and dominant step in 5’SS selection, and the pre-mRNA-snRNA interactions seem to be supported by specific proteins of the spliceosome core. Considering this scenario, strengthened U5 and U6 snRNAs should promote alternative selection of certain 5’SS variants even in the absence of identified B and Bact proteins (see below).

### Alternative selection of certain competing 5’SS is independent of the protein factors

A pre-mRNA variant with upstream 5’SS /GCA<u>U</u>GU (no. 7) was preferentially selected in all mutant strains except for the *prp8-101* allele (Figure 4b and S4b). Since +4U strongly pairs with U6 snRNA’s ACAGA box, these results suggested that the snRNA might have suppressed the selection defects. Non-selection of /GCA<u>U</u>GU in *prp8-101* allele appeared to be specific to the GC donor sites. We explored this possibility by assaying activities from the established *ACT1–CUP1* splicing reporter^38^ with 5’SS variants /GCAAGU, /GCA<u>U</u>GU, and /G<u>U</u>A<u>U</u>GU. Interestingly, /GCA<u>U</u>GU was used efficiently in all mutants, except for *Δhub1* and *prp8-101*. C(+2)U exchanged /G<u>U</u>A<u>U</u>GU made the 5’SS canonical and was efficiently used in all strains, including *Δhub1* and *prp8-101* (Figure 4c). Thus, *prp8-101* allele could not use 5’SS with ‘C’ at +2 position, leading to the loss of alternative splicing. Furthermore, splicing from +3C variant of 5’SS /GU<u>C</u>UGU was defective in *Δhub1*, *Δsnu66 and prp8-101* strains (Figure S4e). Thus, Hub1, Snu66 and Prp8 E1960 surface promote selection of non-canonical 5’SS containing +2C and +3C.

**Figure 4:**
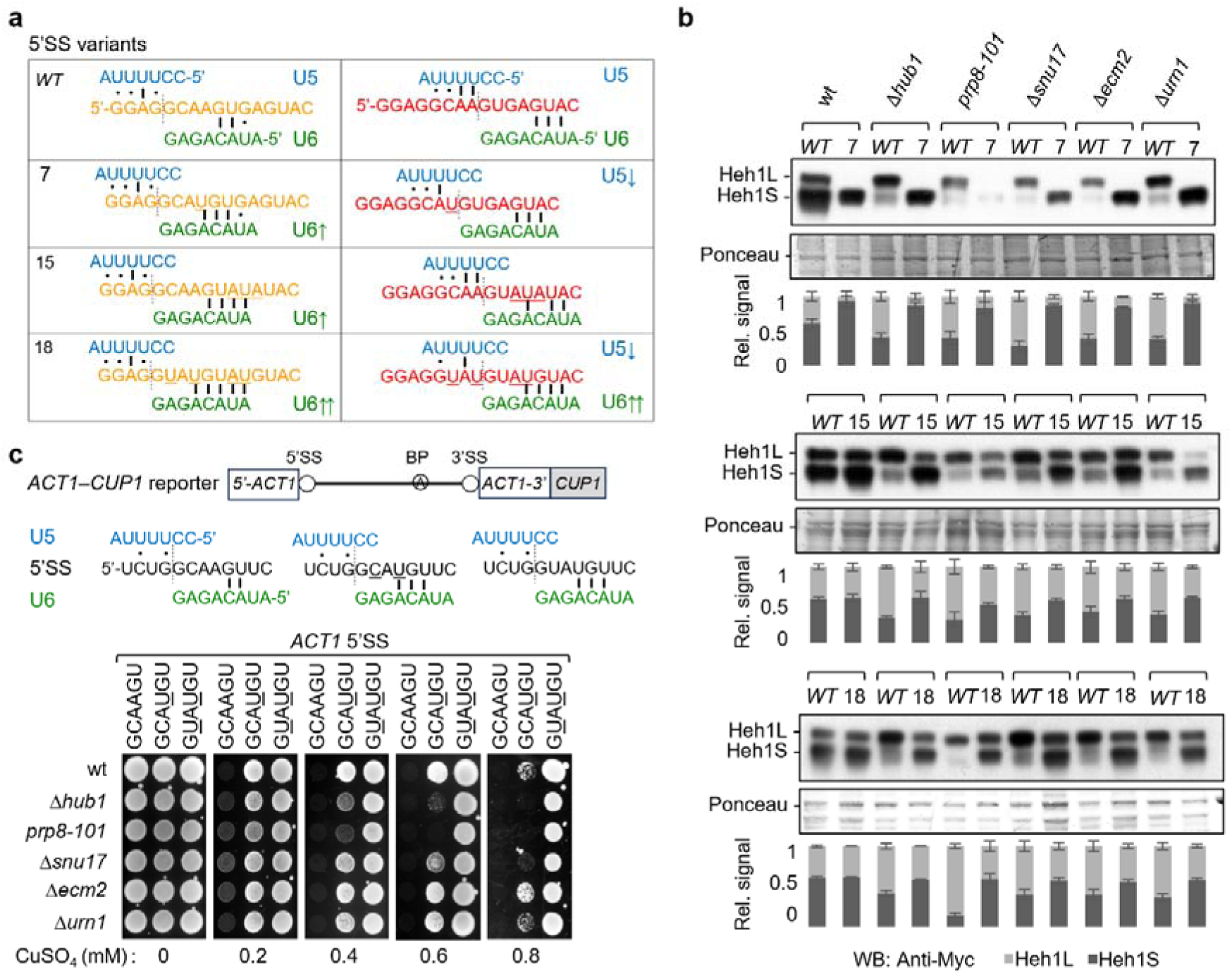
B complex factors are dispensable for alternative splicing from specific 5’SS variants. **(a)** Pairing of U5 and U6 snRNAs to *HEH1 WT* pre-mRNA, variant 7 *HEH1S*•{A(+4)U} or *HEH1L*•{A(−1)U}; variant 15 {*HEH1S*•{G(+7)A,A(+8)U,G(+9)A} or *HEH1L*•{G(+3)A,A(+4)U,G(+5)A}; and variant 18 *HEH1S*•{C(+2)U,A(+4)U,G(+7)A,A(+8)U} or *HEH1L*•{C(−3)U,A(−1)U,G(+3)A,A(+4)U}. **(b)** The effect of variants 7, 15, and 18 on *HEH1* alternative splicing was analyzed in different mutant strains by western blot. Data shown as mean ± s.d. (n=3 independent repeats). **(c)** *ACT1*–*CUP1* reporter was used to understand the outcomes in variant 7. U5 and U6 snRNAs pairing to *HEH1S* 5’SS GCAAGU, GCA<u>U</u>GU and G<u>U</u>A<u>U</u>GU were compared by the reporter assay. The 5’SS mutants were analyzed by growth at indicated CuSO_4_ concentrations, which correlates with their splicing efficiency.

Prp8 is reported to exist in two conformations to maintain equilibrium between the two catalytic steps of splicing. Thus, its opposing alleles rescue splicing defects in one another^39^. However, the defects in the first step allele *prp8-101* could not be explained by this mechanism, since a second step allele *prp8-161* did not restore 5’SS selection in *prp8-101* (Figure S4d).

To find competing 5’SS that would allow alternative splicing through U5 and U6 snRNAs, but without the need of the above-mentioned B and Bact proteins, we made additional pre-mRNA variants by changing more than one nucleotide and monitored their splicing pattern (Figure S2). Proteins expressed from twenty constructs revealed two variants of competing 5’SS that allowed alternative splicing not only in wild-type yeast but also in strains mutated for the B and Bact proteins. 5’SS pairing of U5 and U6 snRNAs was strengthened in the variant 18 containing two canonical 5’SS /G<u>U</u>A<u>U</u>/GU<u>AU</u>GU. The snRNAs’ pairing was recalibrated in variant 15, containing two non-canonical 5’SS /GCAA/GU<u>AUA</u>U. Both these variants allowed alternative splicing not only in cells lacking the protein factors but also in the *prp8-101* allele (Figure 4b and S4b). In variant 18, the competing sites are canonical and are readily used without the need for protein factors. The efficient use of canonical 5’SS in the absence of the protein factors is further supported by *HEH1–CUP1* reporters (Figure S4f-h).

### U5 and U6 snRNAs can promote alternative 5’SS selection without the protein factors

*HEH1S* expression from /GCAA/GU<u>AUA</u>U (variant 15) in cells lacking the protein factors was intriguing, since changes in this variant were beyond the hexanucleotide /GCAAGU. The outcomes could be explained by two possibilities: (i) weakening of the dominant 5’SS /GUGAGU to /GU<u>AUA</u>U rebalanced the competition for the weaker site, or (ii) improved binding of +7U and +8A in /GCAAGU<u>AU</u> to U6 snRNA nucleotides U46 and A45. The data further validated the importance of these two positions of U6 snRNA for alternative splicing discussed in Figure 3.

We tested these possibilities by using two reporters: *HEH1S–CUP1* and *HEH1L–CUP1* (Figure 5b). The protein factors-independent selection of *HEH1S* in /GCAAGU<u>AU</u>, whether due to rebalanced competition or enhanced U6 snRNA pairing, was studied by weakening *HEH1L* 5’SS into /GCAA/GUGAG<u>A</u> (variant 10; lacks potential *HEH1S*-U6 pairing at +7 and +8 positions) and /GCAA/GU<u>AUAA</u> (variant 20; enhances U6 pairing at +7 and +8 positions of *HEH1S*) (Figure 5a and c). 5’SS selection in *Δhub1* and *prp8-101* improved for reporters with enhanced U6 pairing at +7 and +8 positions of the 5’SS, although the defect was only partially restored in the latter. The 5’SS selection in *Δsnu17*, *Δurn1,* and *Δecm2,* on the other hand, was improved by both enhanced U6 snRNA pairing to *HEH1S* and weakening competition from *HEH1L* (Figure 5d). These results suggested that Hub1 and Prp8 promote selection of 5’SS with +2C (Figure 4c), likely by stabilising U6 snRNA (Figure 5d). In addition, Snu17, Urn1, and Ecm2 likely act by stabilising U6 snRNA and also by promoting splicing from the weaker 5’SS present in the vicinity of a dominant one, possibly by improving detection of one another by U5 and U6 snRNAs.

**Figure 5:**
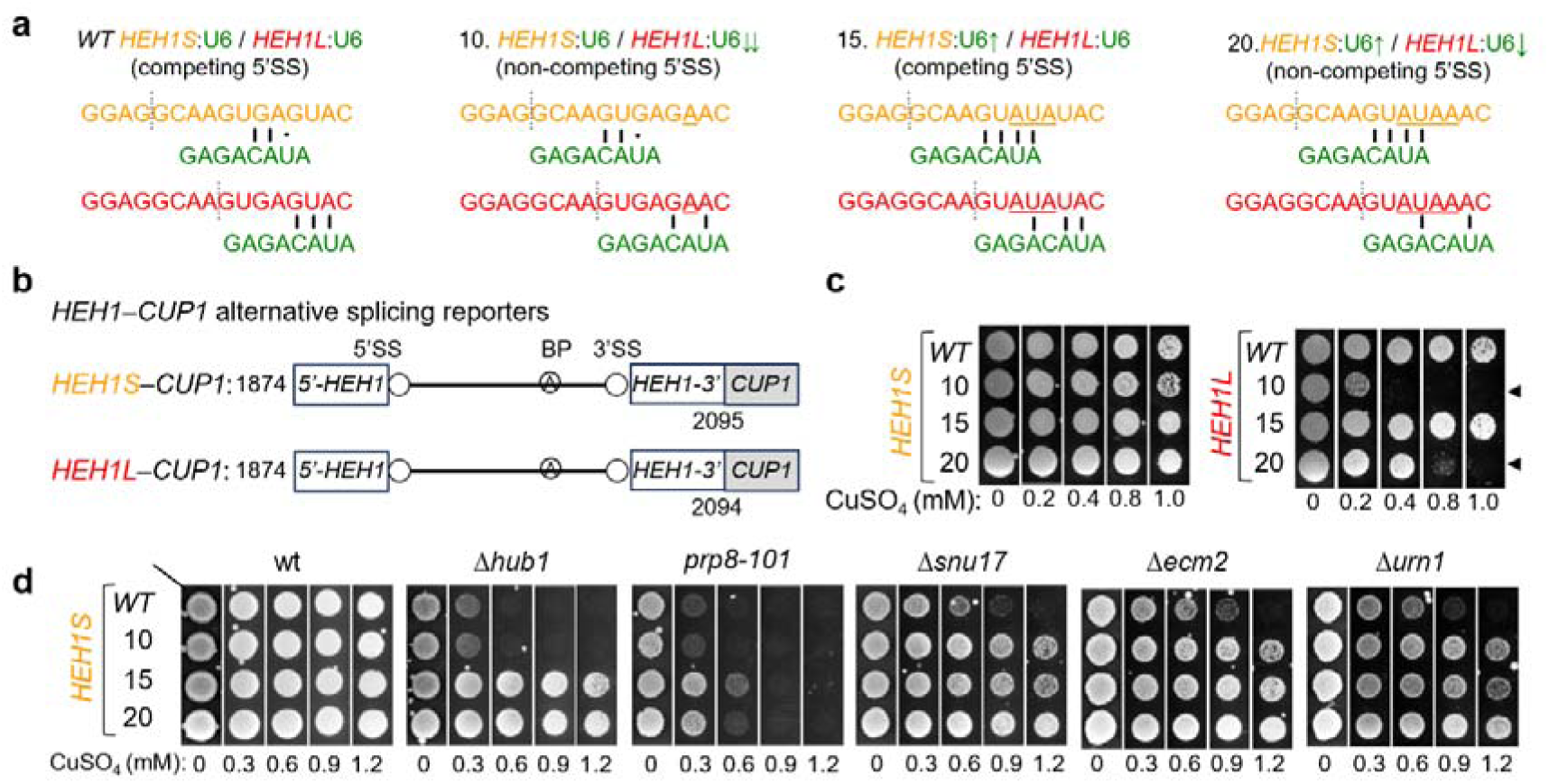
5’SS compete for U6 snRNA. **(a)** Pairing of U6 snRNA with *HEH1 WT* pre-mRNA, variant 10 *HEH1S*•{U(+10)A} or *HEH1L*•{U(+6)A}; variant 15 {*HEH1S*•{G(+7)A,A(+8)U,G(+9)A} or *HEH1L*•{G(+3)A,A(+4)U,G(+5)A}; and variant 20 {*HEH1S*•{G(+7)A,A(+8)U,G(+9)A,U(+10)A} or *HEH1L*•{G(+3)A,A(+4)U,G(+5)A,U(+6)A}. Up/down arrows (↑/↓) indicate increase/decrease in U5 / U6 pairing and competition among the two sites. **(b)** *HEH1*–*CUP1* alternative splicing reporters made by replacing *ACT1* with *HEH1* segment in *ACT1*–*CUP1* reporter. In *HEH1S*–*CUP1, HEH1S* mRNA is in frame with *CUP1*, and in *HEH1L*–*CUP1, HEH1L* mRNA is in frame with *CUP1.* The sequence used was from 1874-2095 for *HEH1S*–*CUP1* and 1874-2094 for *HEH1L*–*CUP1*. **(c)** Reporter activities were tested at different CuSO_4_ concentrations (in mM). (◄) The left pointed triangle denotes the abolished *HEH1L* 5’SS in the variant 10 and weak U6 pairing in the variant 20. **(d)** The *HEH1S*–*CUP1* reporter variants were tested for their splicing efficiency by yeast growth at different CuSO_4_ concentrations.

If the protein factors acted by stabilising U5 and U6 snRNAs on the weaker 5’SS, expression of these snRNA variants with strengthened pairing to the alternative 5’SS should bypass the need for the protein factors. Such U5 and U6 snRNA variants were identified above (Figure 3) for their ability to bind strongly to *HEH1S* 5’SS and promote its competitive selection over the dominant *HEH1L* 5’SS. We tested whether these snRNA variants would also promote a similar preference for *HEH1S* 5’SS in cells lacking the protein factors. Expression of U5 snRNA U(98)C and U6 snRNA U(46)C,A(45)U variants from multicopy plasmids restored alternative splicing defects in *Δhub1*, *Δsnu17*, *Δecm2* and *Δurn1* mutants. The defect was also restored, albeit partially, in the *prp8-101* allele (Figure 6 and Figure S5b,d,e, and f). Despite improved recognition by U6 snRNA, partial rescue of *HEH1S* selection in the *prp8-101* mutants could be due to inefficient catalysis from +2C containing 5’SS (Figure 4c).

**Figure 6:**
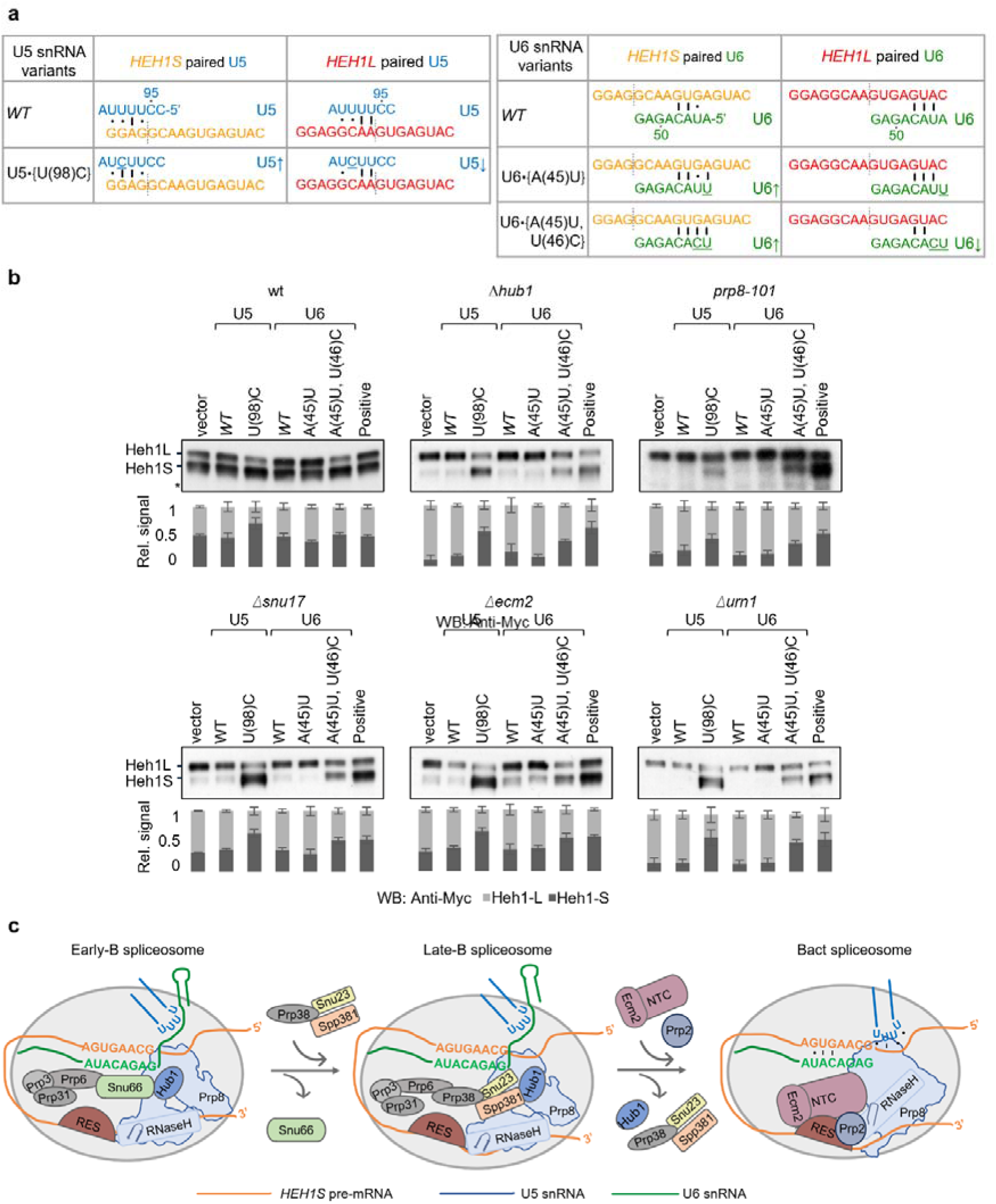
U5 and U6 snRNA variants can bypass the need for B and Bact proteins. **(a)** The pairing of U5•{U(98)C}, U6•{A(45)U and U6•{A(45)U,U(46)C} to *HEH1* alternative isoforms compared to *WT* snRNA. Up / down arrow means (↑/↓) increase/decrease in U5/U6 pairing compared to *WT* snRNA in *HEH1* isoforms. **(b)** Both U5 and U6 variants were overexpressed in different mutant strains for their effect on *HEH1* alternative splicing. The relative abundance of the protein isoforms was monitored by Western blots. Data shown as mean ± s.d. (n=3 independent repeats). **(c)** The schematics show changes from B to Bact complex for the *HEH1* alternative splicing. Snu66 escorts Hub1 to the spliceosome^28^ towards the 5’ exon in early B complex, and hands over to the Spp381 (Prp38 complex subunit) ^53, 56^ in the late B complex. Hub1 interacts with the 5’ exon at the exon-intron boundary. Snu66 and Spp381 binding to the Hub1 may be mutually exclusive. Tri-snRNP factors Prp6, Prp3 and Prp31 are present throughout the B complex. They likely stabilise the U4/U6 snRNA duplex and prevent its premature unbinding^63^. Prp8 changes from a partially open conformation to a closed state from B to Bact^50, 62^. The changes in Prp8 RH domain β-hairpin to loop conformation orchestrate the release of tri-snRNP factors and incorporation of NTC into the spliceosome^48, 62^.

## DISCUSSION

Our findings, along with earlier reports, support a plausible mechanism of alternative 5’SS selection in B and Bact spliceosomes orchestrated by U5 and U6 snRNAs. Hub1 is likely recruited to the early B spliceosome through Snu66 and transferred to the MFAP1 (yeast Spp381) subunit of the Prp38 complex in the late B spliceosome. Core splicing factors centered around Prp8’s RH domain appears to play crucial roles in stabilizing the spliceosome on the alternative GC donor. The RES and NTC proteins promote competition for the alternative 5’SS, possibly from the Bact complex (Figure 6c).

### Pre-mRNA-snRNA interactions in alternative 5’SS selection

The data presented in this study support a dominant role of U5 and U6 snRNAs and a regulatory role of B and Bact proteins in alternative 5’SS selection. Early in the splicing cycle, U1 snRNP is paired at the 5′SS until the pre-B complex. U1 snRNA and associated factors have been reported to promote alternative 5’SS selection^40^. However, we could not find a role of U1 snRNPs and spliceosomes before the pre-B stage in alternative splicing from *HEH1*-type introns. At the B stage of spliceosome assembly, U1 snRNA hands over 5’SS to U6 snRNA. The ACAGA box of U6 base pairs with +3 to +6 nucleotides of the 5’SS. At the same time, U5 snRNA’s loop 1 base pairs with the upstream four nucleotides of the exon^41^.

U5 and U6 snRNAs were essential for alternative 5’SS selection. The defects in the mutants of the B and Bact complexes were suppressed by overexpression of U5 and U6 variants. At the same time, certain pre-mRNA variants with recalibrated pairing to the competing 5’SS were alternatively spliced in the absence of the protein factors. The marking of the target pre-mRNA by these snRNAs in the B complex allows U4/U6 snRNA unwinding and the formation of U2/U6 snRNA pair in the Bact complex^42^. These steps will define the branching site for the first catalytic step for +2C containing 5’SS in the B* complex^43^. Thus, the selection of the alternative 5’SS revolves around their recognition by U5 and U6 snRNAs.

### Specific B and Bact proteins facilitate alternative 5’SS selection

The trans-acting proteins critical for the alternative selection of 5’SS found in this study and reported earlier^28^ belong to the spliceosome B or Bact complexes. Importantly, alternative 5’SS of *HEH1* is the first *in vivo* target of these factors, including the *prp8-101* allele, in *S. cerevisiae*. Splicing from the dominant 5’SS did not require these proteins. The protein factors appear to promote alternative 5’SS selection by stabilizing the spliceosome in a low-fidelity, high-efficiency conformation. These proteins promote alternative 5’SS selection at the expense of the dominant 5’SS. Supporting this notion, Hub1 has been reported to promote error-prone splicing and usage of non-canonical 5’SS^27, 28^. The RES complex proteins Snu17, Bud13, and Urn1 promote non-canonical splicing^44–46^. Ecm2, on the other hand, is known to promote selection of non-canonical as well as competing 5’SS^32^.

*A unique Prp8 allele and its role in 5’SS selection:* Among the 14 distinct Prp8 alleles^47^ tested in this study, the *prp8-101* (E1960K) allele was defective in alternative 5’SS selection. The mutation lies in its RNaseH/RH domain. The Prp8 RH domain toggles between transitional and catalytic conformations^48^. Till the B complex, it remains in a partially closed conformation and promotes fidelity over splicing efficiency. Its first step alleles^49^ and Prp38^50^ have been shown to stabilize the high-fidelity conformation. The defective splicing in *prp8-101* from +2C and +3C containing 5’SS, suggests high-fidelity feature of this mutant and the role of Prp8 in the usage of +2C and +3C containing 5’SS. The RH domain switches to a completely closed conformation in the Bact complex^48, 51^ and positions the RNA for first-step catalysis. Thus, conformational change during B to Bact transition appears to be critical for *HEH1* alternative splicing.

*Prp3, Prp6 and Prp31:* The Role of these proteins was identified through proximity probing with Hub1. These proteins surround the core Prp8 and show a major switch during the conversion of pre-B to B. The 180° shift in the RH domain from pre-B to B, along with tri-snRNP factors, is important for the recognition of alternative 5’SS^52, 53^.

*A new surface of the ubiquitin-like protein Hub1 for 5’SS selection:* Hub1 joins the complex through non-covalent binding to Snu66. The Prp38 complex proteins Prp38, Snu13 and Spp381/MFAP1, associate with the tri-snRNP and help stabilise the B complex. The D22A surface of Hub1 is reported to bind Snu66 across eukaryotes ^28, 54, 55^. The same surface binds MFAP1 in the human B spliceosome^53, 56^. An additional Hub1 surface, N7 and K13, that binds the 5’ exon at -3 and -4 positions in human spliceosomes, was also critical for the alternative 5’SS selection in yeast^56^. D21 of Hub1 also interacts with the same surface of MFAP1^53^. Thus, Hub1 is possibly recruited to the early B complex through HIND domains in Snu66^28^ and may promote alternative 5’SS selection through MFAP1 and RNA interactions in the mature B complex^53, 56^. The RES complex proteins are present during the transition of the B to the Bact^31^.

*The Bact complex*: Brr2 helicase unwinds the U4/U6 duplex, triggering the release of U4 snRNP and the associated proteins (Prp3, Prp6, Prp31, Snu66, Hub1). Prp38 complex is also released at this stage, a key step for full activation of the spliceosome. U6 snRNA forms new interactions with U2 snRNA, assembling the catalytic core. The NTC protein Ecm2 and the RES complex proteins are associated with the Bact complex, highlighting the role of the RES in the remodelling of Bact via Prp2^51^. The RES complex leaves the spliceosome during Bact to B* conversion. All these proteins are important for alternative 5’SS selection.

In conclusion, yeast *HEH1/SRC1* pre-mRNA has proven to be an important tool for understanding alternative splicing through competing and partially overlapping 5’SS. U5 and U6 snRNAs are primary determinants for alternative 5’SS selection. Splicing factors promoting usage of the weak 5’SS reported previously (Hub1, Snu66, Prp38, Prp8), and the ones identified in this study (the B complex proteins Prp3, Prp6, Prp31, and Snu66; the RES complex subunits Bud13, Snu17, Pml1 and Urn1, and the NTC complex (Ecm2), are known to associate with the B or Bact spliceosomes. U5 and U6 snRNAs are essential components of both these assemblies. They pair with target pre-mRNA for alternative 5’SS selection with the help of the above trans-acting proteins.

## LIMITATIONS OF THE STUDY

The study is supported by approaches of molecular biology, biochemistry and yeast genetics. Atomic details will need spliceosome structures during alternative 5’SS selection. *prp8-101,* containing spliceosome assembled on the alternative 5’SS with +2C, will refine the role of the catalytic core.

## RESOURCE AVAILABILITY

A.K.B. or S.K.M. may be contacted for resources used in this study.

## ACKNOWLEDGEMENT

We thank Stefan Jentsch (MPI Biochemistry, Martinsried, Germany) for his support. The work in S.K.M. laboratory has been supported by IISER Mohali and the Centre for Protein Science Design and Engineering (CPSDE) of the Ministry of Human Resource and Development (MHRD), Government of India; the Max Planck Society, Germany; and the Wellcome Trust/DBT India Alliance Fellowship/grant awarded to S.K.M. A.K.B. was supported by ICMR fellowship and IISER Mohali. We acknowledge Anupa T. Anil and B. Mohapatra for their inputs and technical guidance.

## AUTHOR CONTRIBUTION

S.K.M. initiated the study in the Jentsch laboratory at the MPI Biochemistry, Martinsried, Germany. All authors designed and performed the experiments and analysed the data. A.K.B. and S.K.M. collated the data and prepared the manuscript with inputs from all authors.

## STAR METHOD

### KEY RESOURCE TABLE

| Reagent | Resource | Identifier |
| --- | --- | --- |
| Rabbit polyclonal anti-c-myc antibody | Sigma Aldrich | C3956 |
| Rabbit Peroxidase anti-peroxidase soluble antibody | Sigma Aldrich | A0545 |
| Goat anti-rabbit IgG peroxidase antibody | Sigma Aldrich | P1291 |
| High-Capacity cDNA Reverse Transcription Kit | Applied Biosystem | 4368813 |
| NuPAGE™ Bis-Tris Mini Protein Gels, 4–12% | Invitrogen™ | NP0323BOX |
| RQ1 RNase-Free DNase | Promega | M610A |
| Zymo-Spin II Columns | Zymo Research | C1008-250 |
| Vent DNA polymerase | NEB | M0254L |
| **Softwares** |  |  |
| Fiji-windows 64 | NIH | https://imagej.net/software/fiji |
| AlphaFold3 | DeepMind (Google) | https://alphafoldserver.com/ |

### METHOD DETAILS

#### Plasmids, yeast strains, *S. cerevisiae* transformation, and growth assays

Plasmids and strains used in this study are listed in Table 1 and Table 2, respectively. The yJU75 yeast strain, plasmids *PRP8* and *prp8-101,* and *ACT1–CUP1* reporter were kind gifts from C. Guthrie and M. Konarska’s lab. Prp8 mutant strains were made by shuffling out the Ura+ *PRP8* wildtype plasmid with the Trp+ *prp8* mutants on 5-fluoroorotic acid (FOA) plates. *The S. cerevisiae* deletion strains of splicing factors were obtained from the Euroscarf haploid deletion library. Competent cell preparation and transformation were performed following previously published protocols.^57, 58^. Chromosomal tagging for western blot, double knockout strains for genetic interactions, and splicing factors deletion in the yJU75 background were made by using the reported protocol.^57, 58^. Protein overexpression was achieved by expressing clones under a strong *gal promoter*. For growth/spot assays, 5-fold serial dilutions of cells were spotted on the indicated agar plates, and plates were incubated at temperatures indicated in the figure.

**Table 1:** Plasmids used in this study.

| ID | Name | Plasmids | Description | Ref. |
| --- | --- | --- | --- | --- |
| 3658 | WT | YC111 pGAL-3MYC–*HEH1*-tADH | *S. cerevisiae* *HEH1* gene under GAL promoter with 3MYC tag at the N-terminus and ADH terminator | Mishra et al. 2011 |
| D1401 |  | YC111 pGAL-3MYC–*heh1* (A1913T)-tADH | A1913T mutation pGAL 3MYC–*heh1* tADH | This study |
| D1402 |  | YC111 pGAL-3MYC–*heh1* (A1915T)-tADH | A1915T mutation pGAL 3MYC–*heh1* tADH | This study |
| D1403 |  | YC111 pGAL-3MYC–*heh1* (G1916T)-tADH | G1916T mutation pGAL 3MYC–*heh1* tADH | This study |
| D1404 |  | YC111 pGAL-3MYC–*heh1* (G1916A)-tADH | G1916A mutation pGAL 3MYC–*heh1* tADH | This study |
| D1405 |  | YC111 pGAL-3MYC–*heh1* (C1918A)-tADH | C1918A mutation pGAL 3MYC–*heh1* tADH | This study |
| D1406 |  | YC111 pGAL-3MYC–*heh1* (C1918T)-tADH | C1918T mutation pGAL 3MYC–*heh1* tADH | This study |
| D1407 |  | YC111 pGAL-3MYC–*heh1* (A1920T)-tADH | A1920T mutation pGAL 3MYC–*heh1* tADH | This study |
| D1408 |  | YC111 pGAL-3MYC–*heh1* (G1923A)-tADH | G1923A mutation pGAL 3MYC–*heh1* tADH | This study |
| D1409 |  | YC111 pGAL-3MYC–*heh1* (A1924T)-tADH | A1924T mutation pGAL 3MYC–*heh1* tADH | This study |
| D1410 |  | YC111 pGAL-3MYC–*heh1* (T1926A)-tADH | T1926A mutation pGAL 3MYC–*heh1* tADH | This study |
| D1411 |  | YC111 pGAL-3MYC–*heh1* (A1920T, A1924T)-tADH | A1920T, A1924T mutation pGAL 3MYC–*heh1* tADH | This study |
| D1412 |  | YC111 pGAL-3MYC–*heh1* (G1923C, A1924T)-tADH | G1923C, A1924T mutation pGAL 3MYC–*heh1* tADH | This study |
| D1413 |  | YC111 pGAL-3MYC–*heh1* (C1918T, A1919C, A1920T)-tADH | C1918T, A1919C, A1920T mutation pGAL 3MYC–*heh1* tADH | This study |
| D1414 |  | YC111 pGAL-3MYC–*heh1* (A1920T, G1923A, A1924T)-tADH | A1920T, G1923A, A1924T mutation pGAL 3MYC–*heh1* tADH | This study |
| D1415 |  | YC111 pGAL-3MYC–*heh1* (G1923A, A1924T, G1925A)-tADH | G19223A, A1924T, G1925A mutation pGAL 3MYC–*heh1* tADH | This study |
| D1416 |  | YC111 pGAL-3MYC–<br>heh1 (G1923A, A1924T,<br>G1925T, T1926A)-tADH | G1923A, A1924T, G1925T, T1926A<br>mutation pGAL 3MYC–heh1 tADH | This study |
| D1417 |  | YC111 pGAL-3MYC–<br>heh1 (G1923A, A1924T,<br>G1925A, T1926A,<br>A1927C)-tADH | G1923A, A1924T, G1925A, T1926A,<br>A1927C mutation pGAL 3MYC–heh1<br>tADH | This study |
| D1418 |  | YC111 pGAL-3MYC–<br>heh1 (C1918T, A1920T,<br>G1923A, A1924T)-tADH | C1918T, A1920T, G1923A, A1924T<br>mutation pGAL 3MYC–heh1 tADH | This study |
| D1419 |  | YC111 pGAL-3MYC–<br>heh1 (G1921A)-tADH | G1921A mutation pGAL 3MYC–heh1<br>tADH | This study |
| D1420 |  | YC111 pGAL-3MYC–<br>heh1 (G2046C)-tADH | G2046C mutation pGAL 3MYC–heh1<br>tADH | This study |
| D897 |  | YE112 pADH-3myc–<br>Snu66-tADH | *S. cerevisiae* SNU66 gene under<br>GAL promoter with 3MYC tag at the<br>N-terminus and ADH terminator | This study |
| D909 |  | YE112 pADH-3myc–<br>snu66-RRAA-tADH | R16A, R47A mutation pGAL 3MYC–<br>snu66 tADH | This study |
| D605 |  | YC333 pHUB1-Hub1-tADH | *S. cerevisiae* HUB1 gene under its<br>own promoter and terminator | This study |
| D1382 |  | YC333 pHUB1-Hub1(R9A)-tADH | R9A mutation pHUB1-Hub1-tHUB1 | This study |
| D1383 |  | YC333 pHUB1-Hub1(K13A)-tADH | K13A mutation pHUB1-Hub1-tHUB1 | This study |
| D1384 |  | YC333 pHUB1-Hub1(R9A, K13A)-tADH | R9A, K13A mutation pHUB1-Hub1-tHUB1 | This study |
| D739 |  | YC333 pHUB1-Hub1(D22A)-tADH | D22A mutation pHUB1-Hub1-tHUB1 | This study |
| D740 |  | YC333 pHUB1-Hub1(H63L)-tADH | H63L mutation pHUB1-Hub1-tHUB1 | This study |
| 3743 |  | PRP8+ WT TRP CEN | PRP8 with its own promoter and<br>terminator | This study |
| D1421 | *prp8-161* | prp8 (P986T) TRP CEN | prp8 (P986T) with its own promoter<br>and terminator | This study |
| D1422 | *prp8\** | prp8 (P1384L) TRP<br>CEN | prp8 (P1384L) with its own promoter<br>and terminator | This study |
| D1423 | *syf77* | prp8 (L1577F) TRP CEN | prp8 (L1577F) with its own promoter<br>and terminator | This study |
| D1424 | *prp8-107* | prp8 (F1834L) TRP CEN | prp8 (F1834L) with its own promoter<br>and terminator | This study |
| D1425 | *prp8-cat* | prp8 (F1851L) TRP CEN | prp8 (F1851L) with its own promoter<br>and terminator | This study |
| D1426 | *prp8-cat* | prp8 (V1860D) TRP<br>CEN | prp8 (V1860D) with its own promoter<br>and terminator | This study |
| D1427 | *prp8-201* | prp8 (T1861P) TRP CEN | prp8 (T1861P) with its own promoter and terminator | This study |
| D1428 | *prp8-cat* | prp8 (V1862D) TRP CEN | prp8 (V1862D) with its own promoter and terminator | This study |
| D1429 | *prp8-D143* | prp8 (K1864E) TRP CEN | prp8 (K1864E) with its own promoter and terminator | This study |
| D1430 |  | prp8 (T1865K) TRP CEN | prp8 (T1865K) with its own promoter and terminator | This study |
| D1431 | *prp8-162* | prp8 (V1870N) TRP CEN | prp8 (V1870N) with its own promoter and terminator | This study |
| 3745 | *prp8-101* | prp8 (E1960K) TRP CEN | prp8 (E1960K) with its own promoter and terminator |  |
| D1432 | *prp8-153* | prp8 (E1982A) TRP CEN | prp8 (E1982A) with its own promoter and terminator | This study |
| D1433 | *prp8-161, prp8-101* | prp8 (P986T) TRP CEN | prp8 (P986T, E1960K) with its own promoter and terminator | This study |
| D1434 | *prp8-162* | prp8 (V1870N) TRP CEN | prp8 (F1834L, E1960K) with its own promoter and terminator | This study |
| D1435 |  | YC111 pGAL-*SNU17*-tADH | *S. cerevisiae SNU17* gene under GAL promoter and ADH terminator | This study |
| D1436 |  | YC111 pGAL-*ECM2*-tADH | *S. cerevisiae ECM2* gene under GAL promoter and ADH terminator | This study |
| D1437 |  | YC111 pGAL-*URN1*-tADH | *S. cerevisiae URN1* gene under GAL promoter and ADH terminator | This study |
| D1438 | U1•{WT} | pYE195 SNR19-WT + | *S. cerevisiae* SNR19-WT+ gene with its own promoter(1000bp) and terminator(310bp) | This study |
| D1439 | U1•{A7G} | pYE195 snr19 (A7G) | *S. cerevisiae* snr19 (A7G) gene with its own promoter and terminator | This study |
| D1440 | U5•{WT} | pYE195 SNR7-L-WT + | *S. cerevisiae* SNR 7-L-WT+ gene with its own promoter(100bp) and terminator(294bp) | This study |
| D1441 | U5•{U(98)C} | pYE195 snr7-L(U98C) | *S. cerevisiae* snr7-L (U98C) gene with its own promoter and terminator | This study |
| D1442 | U5•{U(98)G} | pYE195 snr7-L(U98G) | *S. cerevisiae* snr7-L (U98G) gene with its own promoter and terminator | This study |
| D1443 | U5•{U(98)A} | pYE195 snr7-L(U98A) | *S. cerevisiae* snr7-L (U98A) gene with its own promoter and terminator | This study |
| D1444 | U6•{WT} | pYE195 SNR6-WT + | *S. cerevisiae* SNR6-WT+ gene with its own promoter(1000bp) and terminator(374bp) | This study |
| D1445 | U6•{A51G} | pYE195 snr6(A51G) | *S. cerevisiae* snr6 (A51G) gene with its own promoter and terminator | This study |
| D1446 | U6•{A51U} | pYE195 snr6(A51U) | *S. cerevisiae* snr6 (A51U) gene with its own promoter and terminator | This study |
| D1447 | U6•{G50U} | pYE195 snr6(G50U) | *S. cerevisiae* snr6 (G50U) gene with its own promoter and terminator | This study |
| D1448 | U6•{G50C} | pYE195 snr6(G50C) | *S. cerevisiae* snr6 (G50C) gene with its own promoter and terminator | This study |
| D1449 | U6•{A49U} | pYE195 snr6(A49U) | *S. cerevisiae* snr6 (A49U) gene with its own promoter and terminator | This study |
| D1450 | U6•{U46C} | pYE195 snr6(U46C) | *S. cerevisiae* snr6 (U46C) gene with its own promoter and terminator | This study |
| D1451 | U6•{A45U} | pYE195 snr6(A45U) | *S. cerevisiae* snr6 (A45U) gene with its own promoter and terminator | This study |
| D1452 | U6•{A45G} | pYE195 snr6(A45G) | *S. cerevisiae* snr6 (A45G) gene with its own promoter and terminator | This study |
| D1453 | U6•{T46C, A45T} | pYE195 snr6(T46C, A45T) | *S. cerevisiae* snr6 (T46C, A45T) gene with its own promoter and terminator | This study |

**Table 2:** Yeast strains used in this study.

| Strain | Genotype | Ref. |
| --- | --- | --- |
| YJU75 (SC11) | *MATa ade2 cup1Δ::ura3 his3 leu2 lys2 prp8Δ::LYS2 trp1; pJU169 (PRP8 URA3 CEN ARS)* | Gift from C. Guthrie |
| SC87 | YJU75 *hub1::KanMX6* | This study |
| SC196 | YJU75 PRP8 TRP | This study |
| SC197 | YJU75 *prp8-101 TRP* | This study |
| SC198 | YJU75 PRP8 TRP *hub1::KanMX6* | This study |
| SC200 | YJU75 *P<sub>CUP1-1</sub>TAP-SRC1::natNT2* | This study |
| SC214 | YJU75 *prp8-101P<sub>CUP1-1</sub>TAP-SRC1::natNT2* | This study |
| SC224 | YJU75 *hub1::KanMX6 P<sub>CUP1-1</sub>TAP-SRC1::natNT2* | This study |
| SC297 | SC200 *nam8::KanMX6* | This study |
| SC321 | SC11. *Dib1-hub1(D22A):: natNT2* | This study |
| SC322 | SC11. *Prp31-hub1(D22A):: natNT2* | This study |
| SC323 | SC11. *Sad1-hub1(D22A):: natNT2* | This study |
| SC324 | SC87. *Dib1-hub1(D22A):: natNT2* | This study |
| SC325 | SC87. *Prp31-hub1(D22A):: natNT2* | This study |
| SC326 | SC87. *Sad1-hub1(D22A):: natNT2* | This study |
| BY4741 | *MATa his3Δ1 leu2Δ0 met15Δ0 ura3Δ0* | This study |
| YSKM601 | BY4741 *hub1 ::KanMX6 P<sub>CUP1-1</sub>TAP-SRC1::natNT2* | This study |
| YSKM602 | BY4741 *lea1::KanMX6 P<sub>CUP1-1</sub>TAP-SRC1::natNT2* | This study |
| YSKM603 | BY4741 *ecm2::KanMX6 P<sub>CUP1-1</sub>TAP-SRC1::natNT2* | This study |
| YSKM604 | BY4741 *bud13::KanMX6 P<sub>CUP1-1</sub>TAP-SRC1::natNT2* | This study |
| YSKM605 | BY4741 *nup60::KanMX6 P<sub>CUP1-1</sub>TAP-SRC1::natNT2* | This study |
| YSKM606 | BY4741 *lsm1::KanMX6 P<sub>CUP1-1</sub>TAP-SRC1::natNT2* | This study |
| YSKM607 | BY4741 *sky1::KanMX6 P<sub>CUP1-1</sub>TAP-SRC1::natNT2* | This study |
| YSKM608 | BY4741 *snu17::KanMX6 P<sub>CUP1-1</sub>TAP-SRC1::natNT2* | This study |
| YSKM609 | BY4741 *pml1::KanMX6 P<sub>CUP1-1</sub>TAP-SRC1::natNT2* | This study |
| YSKM610 | BY4741 *urn1::KanMX6 P<sub>CUP1-1</sub>TAP-SRC1::natNT2* | This study |
| YSKM611 | BY4741 *mlp2::KanMX6 P<sub>CUP1-1</sub>TAP-SRC1::natNT2* | This study |
| YSKM612 | BY4741 *mlp1::KanMX6 P<sub>CUP1-1</sub>TAP-SRC1::natNT2* | This study |
| YSKM613 | BY4741 *pml39::KanMX6 P<sub>CUP1-1</sub>TAP-SRC1::natNT2* | This study |
| YSKM614 | BY4741 *brr1::KanMX6 P<sub>CUP1-1</sub>TAP-SRC1::natNT2* | This study |
| YSKM615 | BY4741 *mud2::KanMX6 P<sub>CUP1-1</sub>TAP-SRC1::natNT2* | This study |
| YSKM616 | BY4741 *mud1::KanMX6 P<sub>CUP1-1</sub>TAP-SRC1::natNT2* | This study |
| YSKM617 | BY4741 *prp17::KanMX6 P<sub>CUP1-1</sub>TAP-SRC1::natNT2* | This study |
| YSKM618 | *BY4741 npl3::KanMX6 P<sub>CUP1-1</sub>TAP-SRC1::natNT2* | This study |
| YSKM619 | *BY4741 cwc15::KanMX6 P<sub>CUP1-1</sub>TAP-SRC1::natNT2* | This study |
| YSKM620 | *BY4741 cwc21::KanMX6 P<sub>CUP1-1</sub>TAP-SRC1::natNT2* | This study |
| YSKM621 | *BY4741 cwc27::KanMX6 P<sub>CUP1-1</sub>TAP-SRC1::natNT2* | This study |
| YSKM622 | *BY4741 ntc20::KanMX6 P<sub>CUP1-1</sub>TAP-SRC1::natNT2* | This study |
| YSKM623 | *BY4741 ntc30::KanMX6 P<sub>CUP1-1</sub>TAP-SRC1::natNT2* | This study |
| YSKM624 | *BY4741 ntc31::KanMX6 P<sub>CUP1-1</sub>TAP-SRC1::natNT2* | This study |
| W303 | *ade2-1 his3-11, 15 leu2-3, 112 ura3 trp1-1 ssd1 can1-100* | Gift from K. Nasmyth |
| YSKM625 | *W303a P<sub>CUP1-1</sub>TAP-SRC1::KanMX6* |  |
| YSKM626 | *W303a P<sub>CUP1-1</sub>TAP-SRC1::KAN Prp8-hub1(D22A)* | This study |
| YSKM627 | *W303a P<sub>CUP1-1</sub>TAP-SRC1::KAN Snu66-hub1(D22A)* | This study |
| YSKM628 | *W303a P<sub>CUP1-1</sub>TAP-SRC1::KAN Prp38-hub1(D22A)* | This study |
| YSKM629 | *W303a P<sub>CUP1-1</sub>TAP-SRC1::KAN Snu66-hub1(D22A) Prp38-hub1(D22A)* | This study |
| YSKM630 | *W303a hub1::HIS3MX6 P<sub>CUP1-1</sub>TAP-SRC1::KanMX6 SNU17-hub1(D22A):: natNT2* | This study |
| YSKM631 | *W303a hub1::HIS3MX6 P<sub>CUP1-1</sub>TAP-SRC1::KanMX6 PRP4-hub1(D22A):: natNT2* | This study |
| YSKM632 | *W303a hub1::HIS3MX6 P<sub>CUP1-1</sub>TAP-SRC1::KanMX6 BRR2-hub1(D22A):: natNT2* | This study |
| YSKM633 | *W303a hub1::HIS3MX6 P<sub>CUP1-1</sub>TAP-SRC1::KanMX6 PRP6-hub1(D22A):: natNT2* | This study |
| YSKM634 | *W303a hub1::HIS3MX6 P<sub>CUP1-1</sub>TAP-SRC1::KanMX6 PRP3-hub1(D22A):: natNT2* | This study |
| YSKM635 | *W303a hub1::HIS3MX6 P<sub>CUP1-1</sub>TAP-SRC1::KanMX6 LEA1-hub1(D22A):: natNT2* | This study |
| YSKM636 | *W303a hub1::HIS3MX6 P<sub>CUP1-1</sub>TAP-SRC1::KanMX6 PRP9-hub1(D22A):: natNT2* | This study |
| YSKM637 | *W303a hub1::HIS3MX6 P<sub>CUP1-1</sub>TAP-SRC1::KanMX6 PRP11-hub1(D22A):: natNT2* | This study |
| YSKM638 | *W303a hub1::HIS3MX6 P<sub>CUP1-1</sub>TAP-SRC1::KanMX6 MUD1-hub1(D22A):: natNT2* | This study |
| YSKM639 | *W303a hub1::HIS3MX6 P<sub>CUP1-1</sub>TAP-SRC1::KanMX6 NAM8-hub1(D22A):: natNT2* | This study |
| SC186 | *W303a snu66::HIS3MX6* | This study |

#### Chromosomal fusion of *hub1*–*D22A* to the C-termini of splicing factors

Splicing by Overlap Extension (SOE) PCR was done to make *hub1(D22A)* C-terminal fusion to essential splicing factors. This fusion was done based on the published protocol^57^. Three sets of primers were used to amplify three fragments. First set of primers amplified a 200-300 bp fragment of the C termini of the desired gene, excluding the stop codon. The reverse primer of this had an overhang of 20-25 bps that was complementary to the 20-25 bps of *hub1(D22A)* N-terminus. The second set of primers amplified the full-length *hub1(D22A)* with the Nat antibiotic selection marker cassette from the plasmid clones of pFA6a-*hub1(D22A)- natNT2*. The third set of primers amplified a fragment of 200-300 bp from the stop codon of the gene that was going to be tagged. The forward primer for this had an overhang of 20-25 bp with the end of the antibiotic-resistance marker. All three fragments were amplified using Vent DNA polymerase. These fragments were confirmed based on their size on the agarose gel and the correct fragments were gel-purified. All three fragments were mixed and joined by SOE PCR using Vent DNA polymerase using the forward primer of the first fragment and the reverse primer of the third fragment. The joined fragment was confirmed by agarose gel electrophoresis and transformed into the *S. cerevisiae* strain. The transformants were selected on agar plates with antibiotic. The chromosomal fusion was confirmed by colony PCR using a forward primer specific to the gene and a reverse primer specific to the antibiotic cassette.

#### Splicing reporters

The splicing reporters used in this study are listed in Table 3. Splicing reporters are modified forms of the conventional *ACT1–CUP1*^38^ and *RP51–LACZ*^30^. The *HEH1S–CUP1 reporters have a 121 bp segment of the S. cerevisiae HEH1* gene (1874-2095 nucleotides), including the intron, using BamHI and KpnI restriction sites. The *HEH1S* mRNA is spliced using GCAAGU 5’SS, generating in-frame *CUP1* mRNA for translation. The methionine initiation codon was introduced by inserting a point mutation at C1881G. Initiation codon and variants of the 5’SS were made by site-directed mutagenesis (SDM) using specific primers, and the change was confirmed using Sanger sequencing. For *HEH1L–CUP1* reporter, a 120 bp segment of the *HEH1* gene (1874-2094 nucleotides) was used to bring the *HEH1L* mRNA spliced using GUGAGU 5’SS in-frame with the *CUP1* gene. For reporter assays, competent cells from the *S. cerevisiae* strain yJU75 were transformed with the reporters, and transformants were selected in media lacking leucine. 1 OD_600_ cells were spotted on solid media with different concentrations of CuSO_4_. The plates were incubated for 2-3 days at 30°C. Similarly, for *HEH1–LACZ* reporters, the *RP51* part of the *RP51–LACZ* reporter (gift from M. Rosbash) was replaced with a *HEH1* fragment of 399 bp (1792-2191 nucleotides) and 398 bp (1792-2190 nucleotides) to obtain *HEH1S–LACZ* and *HEH1L–LACZ* reporters, respectively.

**Table 3:** Splicing reporters used in this study.

| Splicing Reporters | Description |
| --- | --- |
| *ACT1-CUP1* reporter | i) 5'SS GUAUGU (Gift from C. Guthrie)<br>ii) 5'SS GCAUGU (D1454)<br>iii) 5'SS GCAAGU (D1455) |
| *HEH1L-CUP1* reporter | *ACT1* exon1-intron-exon2 in *ACT1-CUP1* reporter is replaced with *HEH1* exon1-intron-exon2 (nucleotides 1874-2094) to keep *HEH1L* 5'SS GUGAGU in frame with *CUP1*. Its variants were made using SDM.<br>i) 5'SS <u>GUGAGU</u>AC (D1456)<br>ii) 5'SS <u>GUGAGA</u>AC (D1457)<br>iii) 5'SS <u>GUAUAU</u>AC (D1458)<br>iv) 5'SS <u>GUAUAA</u>AC (D1459)<br>v) 5'SS <u>GUAUGU</u>AC (D1460) |
| *HEH1S-CUP1* reporter | *ACT1* exon1-intron-exon2 in *ACT1-CUP1* reporter is replaced with *HEH1* exon1-intron-exon2 (nucleotide 1874-2095) to keep *HEH1S* 5'SS GCAAGU in frame with *CUP1*. Its variants were made using SDM.<br>i) 5'SS <u>GCAAGU</u>GUGAGUAC (D1461)<br>ii) 5'SS <u>GCAAGU</u>GUGAGAAC (D1462)<br>iii) 5'SS <u>GCAACU</u>GUAUAUAC (D1463)<br>iv) 5'SS <u>GCAAGU</u>GUAUAAAC (D1464)<br>v) 5'SS <u>GCAAGU</u>GUAUGUAC (D1465) |
| *HEH1L-LACZ* reporter | *RP51* exon1-intron-exon2 is replaced with *HEH1* exon1-intron-exon2 (nucleotide 1792-2190) is fused with the *LACZ* gene to keep *HEH1L* 5'SS GUGAGU in frame with *LACZ*. |
| *HEH1S-LACZ* reporter | *RP51* exon1-intron-exon2 is replaced with *HEH1* exon1-intron-exon2 (nucleotide 1792-2191) is fused with the *LACZ* gene to keep *HEH1S* 5'SS GCAAGU in frame with *LACZ*. |

#### **β**-galactosidase assay

β-galactosidase activity was observed on both solid and liquid media. The X-gal overlay and the ONPG assay were performed by following published protocols^59^.

#### RNA isolation and cDNA synthesis

RNA isolation and cDNA synthesis were done as described previously^60^. Briefly, five OD_600_ cells in the log phase were harvested. Total RNA was isolated by the hot acid phenol method using 15 mL phase lock tubes for phase separation. Residual DNA was removed by treating RNA with DNase I for 15 min at room temperature, followed by RNA clean-up using a Zymo-Spin^TM^ II column. cDNA synthesis from 2μg total RNA was done using reverse transcriptase (RT) and random-hexamer primers at 37°C for 2h. Splicing defects were monitored by detecting intron-containing transcripts or post-splicing mature transcripts using exonic primers across the intron and analysed by agarose gel electrophoresis. For cDNA sequencing, five identical PCR reactions were performed. The PCR products were pooled and kept overnight for precipitation after adding 2.5 times the volume of isopropanol and 1/10^th^ the volume of 3M sodium acetate at -20°C. DNA was pelleted by centrifugation for 15 min at 4°C at maximum speed. The pellet was washed twice with 70% ethanol. Dried pellet was dissolved in 30μl of nuclease-free water, and 10μl of the DNA was sequenced by Sanger sequencing using a primer nested inside the forward primer used in amplification.

#### Quantification of *HEH1* mRNA isoforms

*HEH1* alternative splicing was analysed using RT-PCR followed by cDNA sequencing. *HEH1S* and *HEH1L* mRNAs differ in GCAA nucleotides at their exon junctions (*HEHL* gains them due to the usage of downstream 5’SS), the electropherogram shows mixed peaks corresponding to the two isoforms after the exon-exon junction. The area under the peaks was integrated to find the relative abundance of the two mRNAs. Both common and isoform-specific peaks were analysed within the region corresponding to the 5’exon/intron boundary. To understand alternative 5’SS usage, common nucleotides after the junction in the two isoforms was excluded from further analysis. For each site, we recorded the nucleotide proportions possible for both the cDNAs and calculated the ratio between the two nucleotides, normalising their combined total to 100%. The average value of both forms is directly related to their respective 5’SS usage.

Relative expression of *HEH1L* in wild-type and yeast mutants was obtained from the cDNA electropherogram. The areas under 10 common and 10 mixed peaks (before and after the junction, respectively), were integrated and averaged. Similarly, an area under the peaks specific to *HEH1L* was integrated and averaged. The ratio of common and *HEHL* peaks for different strains was obtained. The fold difference in *HEH1L* expression in the mutants was calculated against the wildtype strain by dividing the average of the area under *HEH1L* peaks (after the junction) by the average of the area under common peaks (before the junction).

#### Western blots

For protein western blots (immunoblotting), logarithmically growing cells equivalent to 2 OD_600_ were harvested. Whole protein was extracted using the trichloroacetic acid (TCA) method^57^. Total proteins used for western blots were denatured by heating cells in a high urea buffer at 65°C for 15 min and then centrifuged. The soluble protein was separated on sodium dodecyl sulphate-polyacrylamide gel electrophoresis (SDS-PAGE) 8% gels, or NuPAGE^TM^ 4%-12% Bis-Tris gels (Invitrogen), transferred to a polyvinylidene difluoride (PVDF) membrane, and probed with specific primary and secondary antibodies. Protein bands were quantified using ImageJ following the published method^61^.

## Supplementary Figures

**Figure S1:**
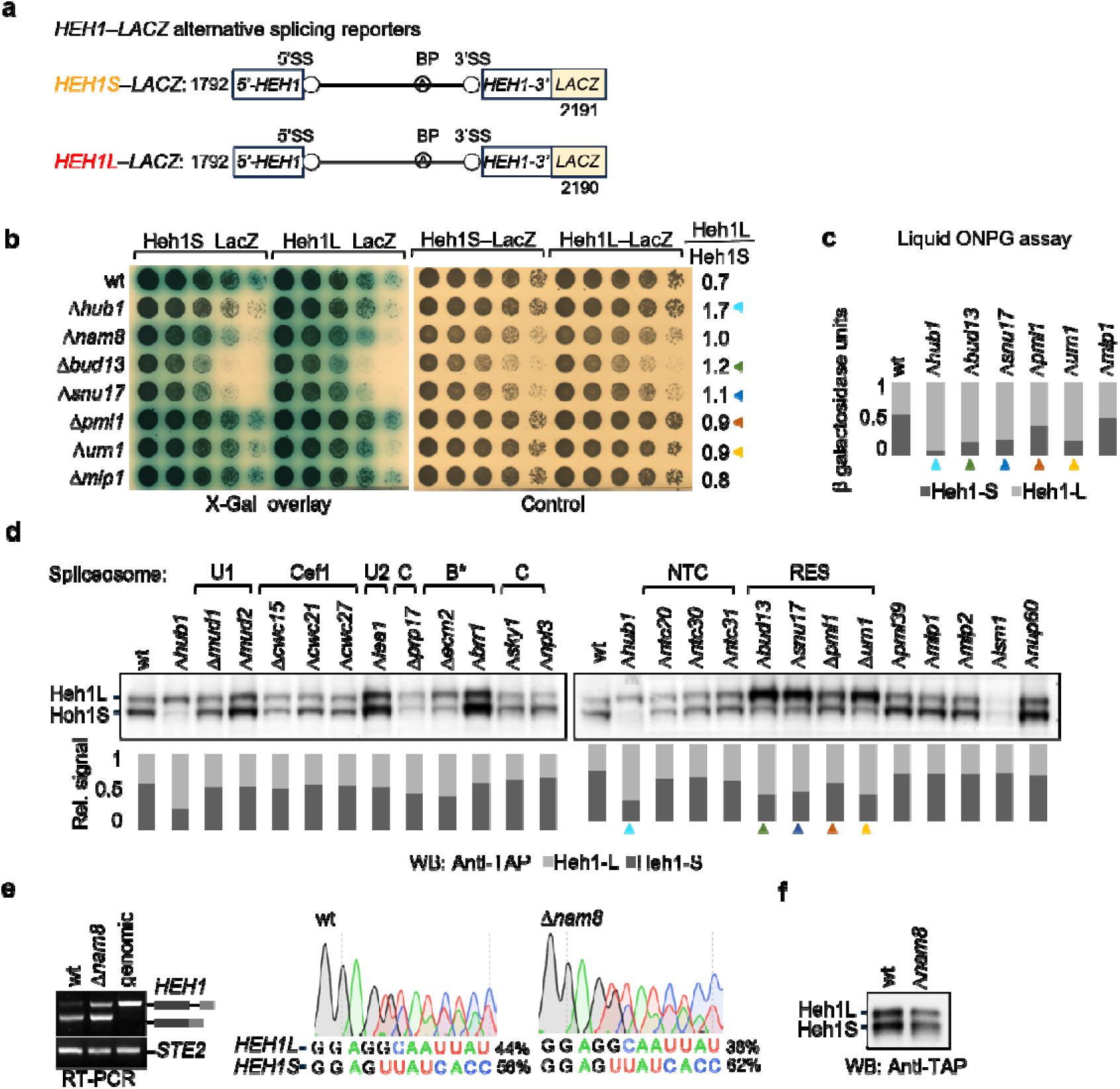
*HEH1* alternative splicing factors. **(a-f)** Screening of yeast deletion strains to identify trans-acting proteins for *HEH1* alternative splicing. **(a)** Schematic showing the *HEH1*–*LACZ* alternative splicing reporters. Reporters modified from the *RP51*–*LACZ* reporter^1^ and designed in a way to specifically monitor the usage of *HEH1S* and *HEH1L* 5’SS. 1 nucleotide is less in *HEH1L*–*LACZ* to bring *HEH1L* 5’SS in frame with *LACZ*. **(b)** β-galactosidase activity was observed on both solid and liquid media. Spot assay of the transformants with splicing reporters was performed on the selection marker plate and overlayed with X-gal. The intensity of blue color, indicative of β-galactosidase activity, correlated with the splicing efficiency. **(c)** Liquid ONPG assay for the indicated strains with splicing reporters. **(d)** Heh1 protein isoforms from N-terminally TAP-tagged *HEH1* in deletion mutants of splicing factors representative of different spliceosomal complexes were monitored by western blots. RT-PCR followed by cDNA sequencing **(e)** and western blot **(f)** showed normal *HEH1* mRNA and protein isoforms in *Δnam8*, thus ruling out Nam8’s (a U1snRNP component) in alternative 5’SS selection. Triangles (▴) with color codes indicate alternative splicing defect observed in different stains in figures b, c, and d.

**Figure S1:**
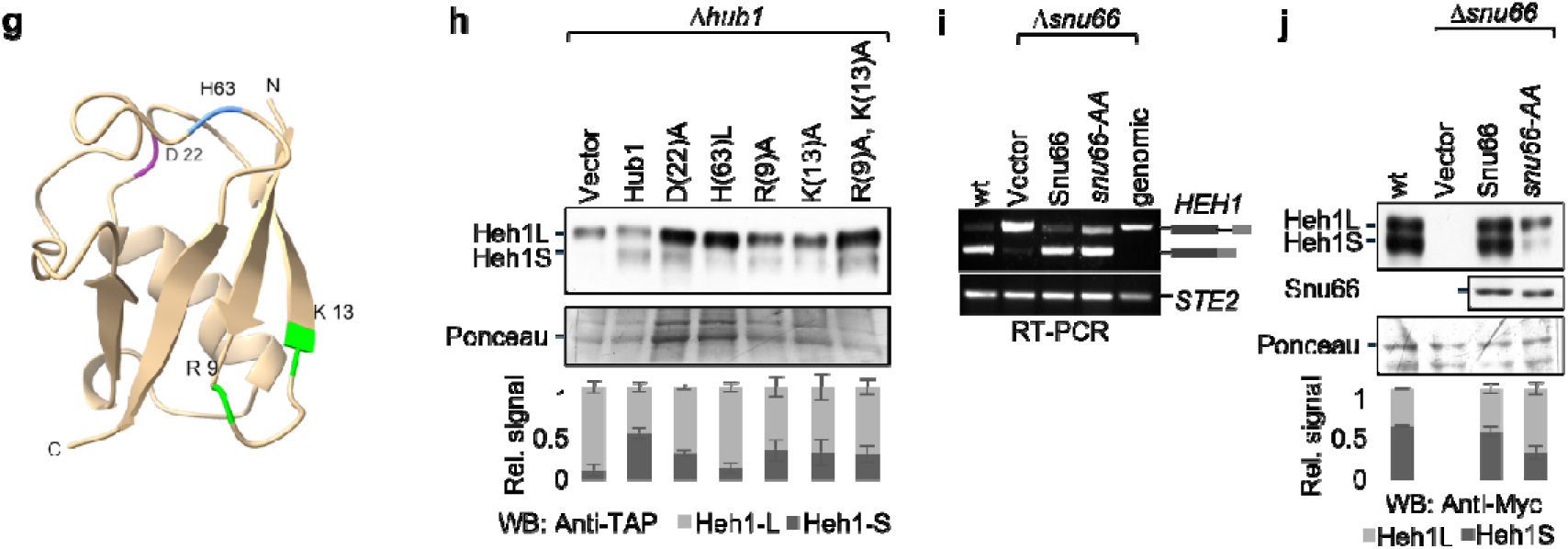
*HEH1* alternative splicing factors. **(g-j)** Hub1 and Snu66 surfaces are involved in *HEH1* alternative splicing. **(g)** AlphaFold predicted structure of yeast Hub1 highlighting its Snu66 binding surface D22, Prp5 binding surface H63, and its presumed RNA binding surfaces R9 and K13. **(h)** *Δ hub*1 strain was transformed with the indicated Hub1 mutants, and Heh1 protein was examined by western blot. Data shown as mean ± s.d. (n=3 independent repeats). **(i)** RT-PCR and **(j)** western blot analysis of wild type (wt), *Δsnu66,* and *snu66-AA* strains. *snu66-AA* refers to Hub1 binding deficient R16A and R47A variant. Data shown as mean ± s.d. (n=3 independent repeats).

**Figure S1:**
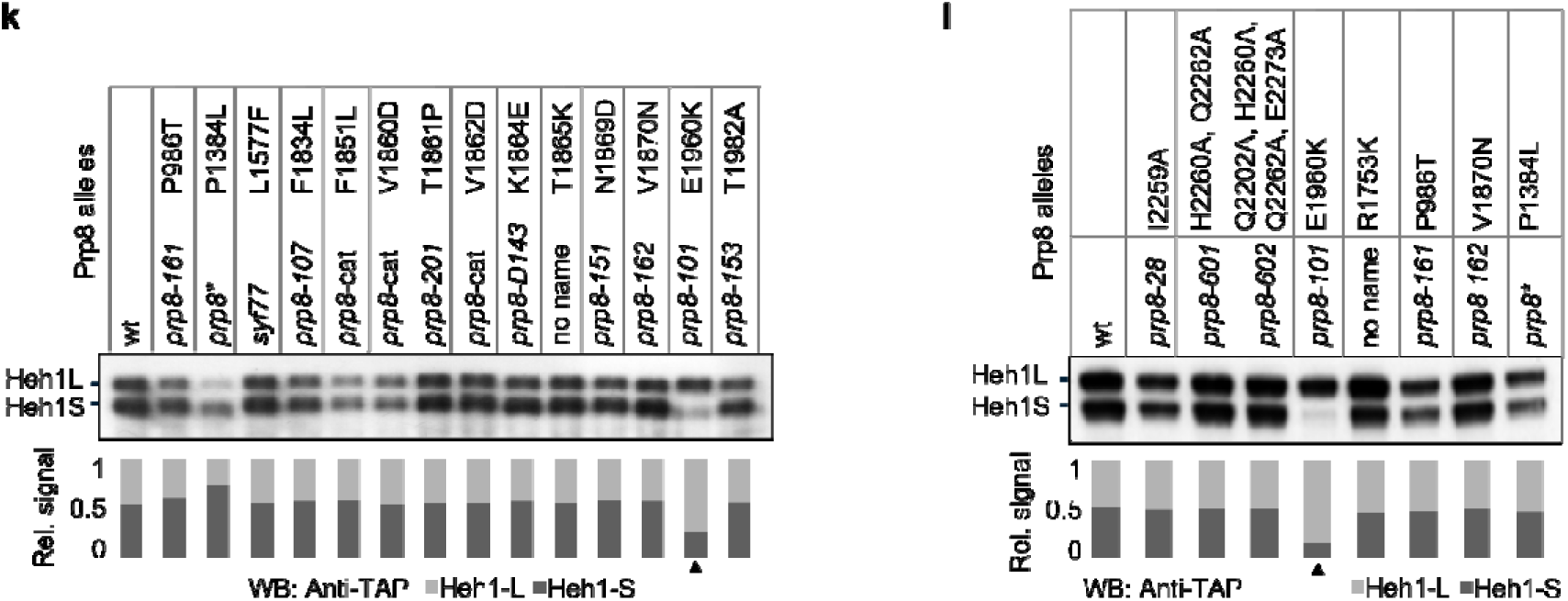
*HEH1* alternative splicing factors. **(k-l)** Screening of Prp8 alleles for defects in *HEH1* alternative splicing. Different alleles of the core splicing factor Prp8 were screened by western blot for the *HEH1* alternative splicing defect. The upward triangle (▲) points at the allele showing the defect.

**Figure S1:**
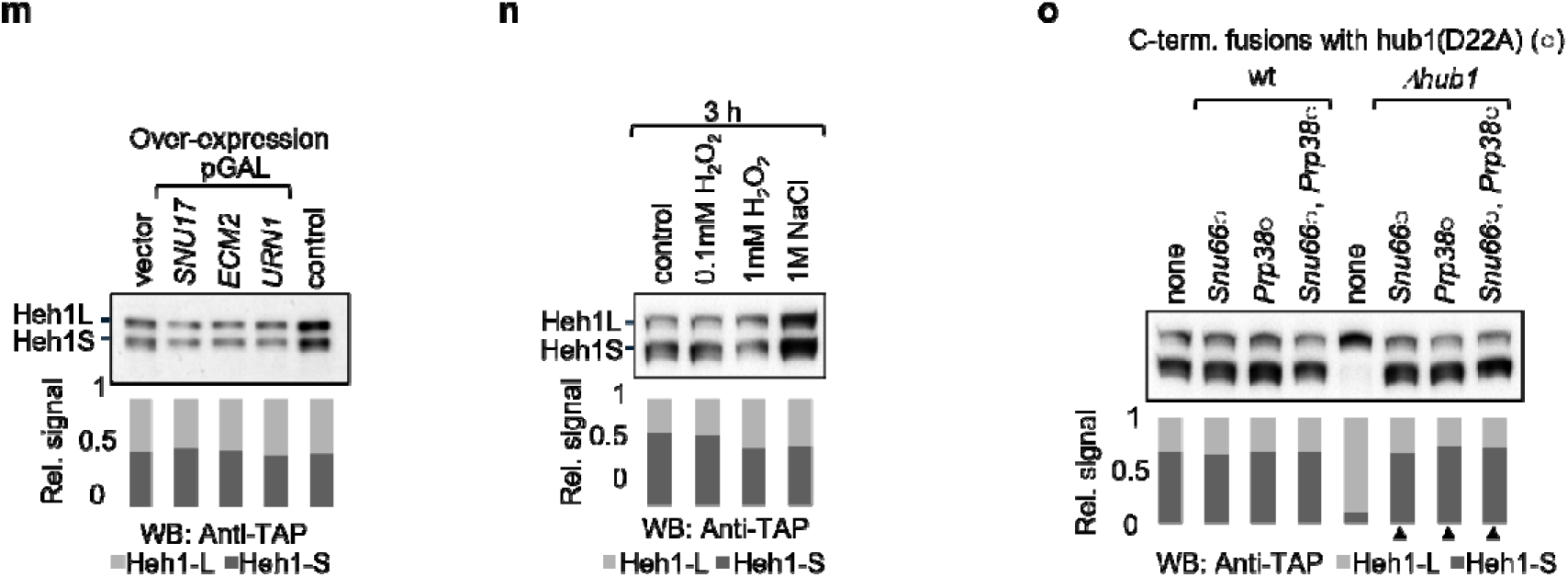
*HEH1* alternative splicing factors. **(m-o)** Overexpression of the trans-acting factor does not alter *HEH1* 5’SS selection. **(m)** *SNU17*, *ECM2,* and *URN1* were expressed in the *wt* strain. Transformants were induced for 5-6 h in 1% galactose media, harvested, and tested for their effect on *HEH1* alternative splicing. (**n)** The effect of stress-induced upregulation of Hub1 on *HEH1* alternative splicing was tested by western blot. **(o)** indicated factors screened by Western blot in the wt and *Δhub1* strains. ○ indicates C-terminal fusion of *hub1(D22A)* with the splicing factors. Upward triangles (▲) indicate proteins that rescued *HEH1* alternative splicing

**Figure S2:**
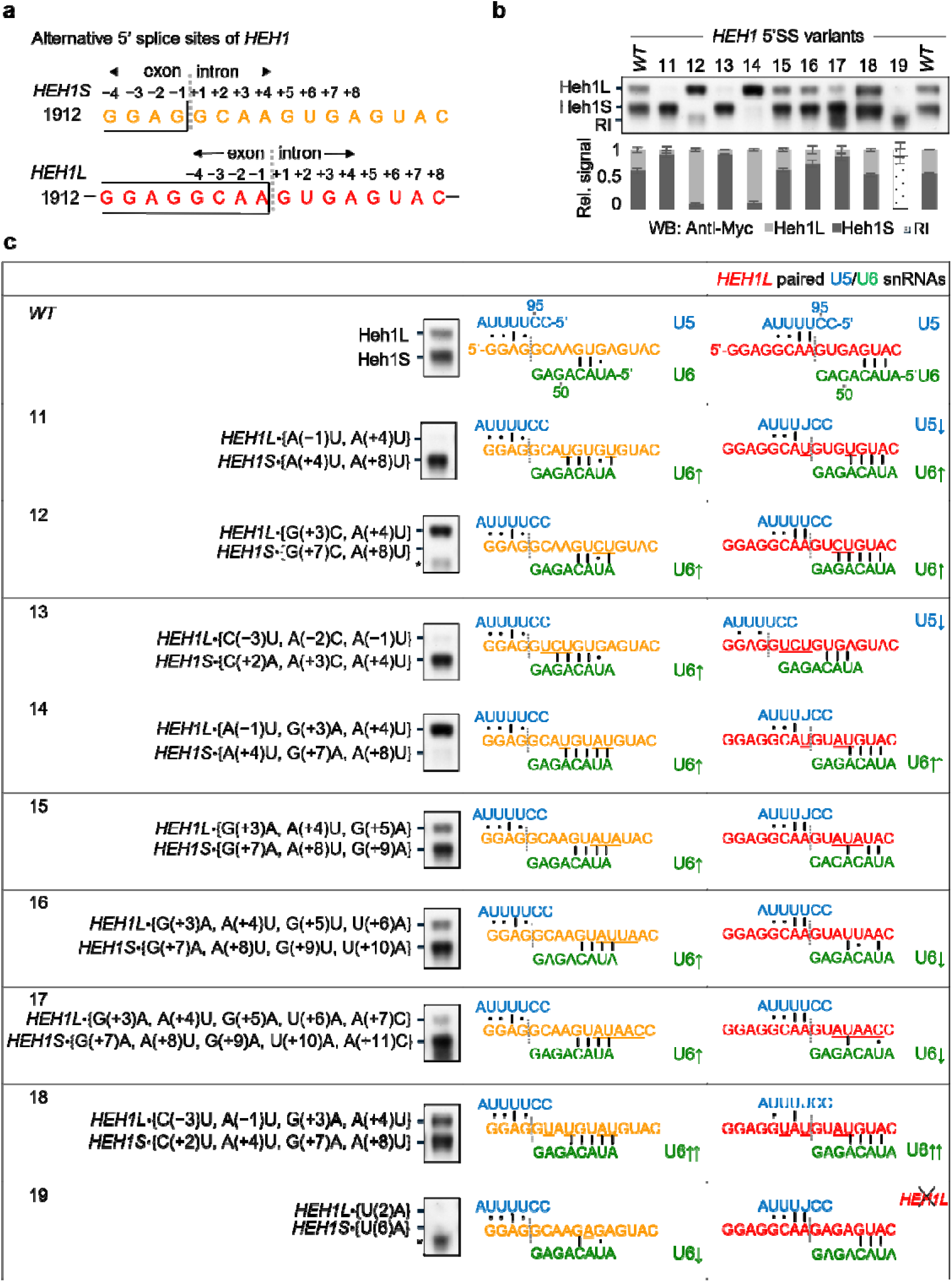
Pairing of pre-mRNA variants with U5 and U6 snRNA and the outcome of *HEH1* alternative splicing. **(a-c)** Sequence at the exon-intron boundary determines 5’SS selection. **(a)** The 16-nucleotide segment at the exon/intron (1913-1926) boundary of *HEH1* was investigated for alternative splicing. The sequence was selected based on reporter assays described earlier, pairing of U5 snRNA loop1 and U6 snRNA ACAGA box. *HEH1S* (orange color) and *HEH1L* (red color) sequences represent isoforms of *HEH1* pre-mRNA. **(b)** More than one mutation was introduced based on U5 and U6 snRNA binding strength to both the 5’SS. The impact of the U5 and U6 interaction change on *HEH1* alternative splicing was monitored in western blots. Data shown as mean ± s.d. (n=3 independent repeats). RI (retained intron) represents the *HEH1* pre-mRNA band. **(c)** Pre-mRNA variants with respective U5 snRNA (blue colour) and U6 snRNA (green colour) pairing. The mutants were analyzed for presumed U5 and U6 snRNA pairings, adapted from^2, 3^. x indicates no *HEH1L* protein in the variant. Western blot lanes are taken from (b). Labels are the same as in Figure 1.

**Figure S2:**
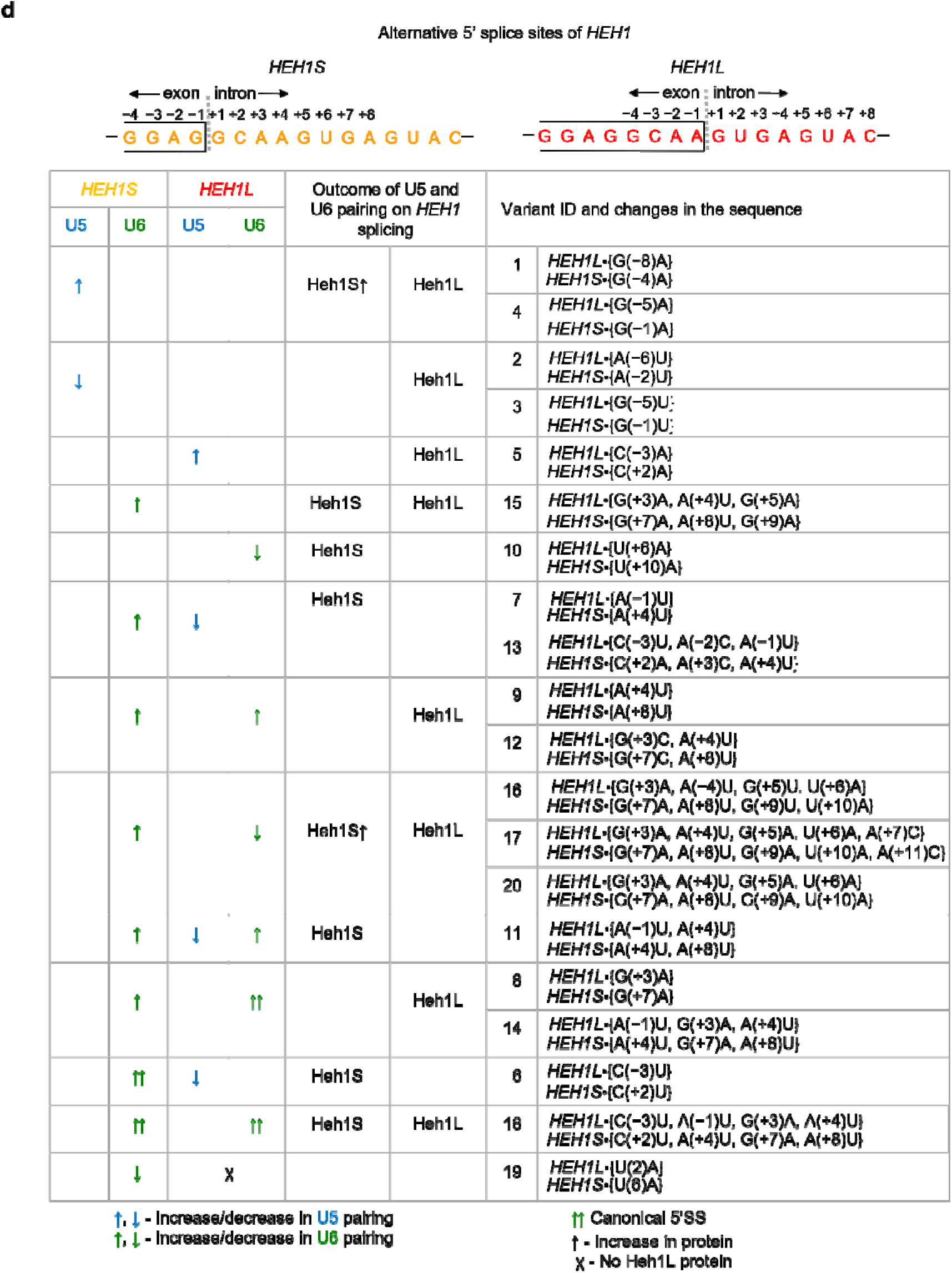
Pairing of pre-mRNA variants with U5 and U6 snRNA and the outcome of *HEH1* alternative splicing. **(d)** Sequence at the exon-intron boundary determines 5’SS selection**(d)** Summarizes how the pairing of U5 and U6 snRNA to the alternative 5’SS decide the splicing outcome of *HEH1* based on the data from Figure 1 and Figure S1.

**Figure S3:**
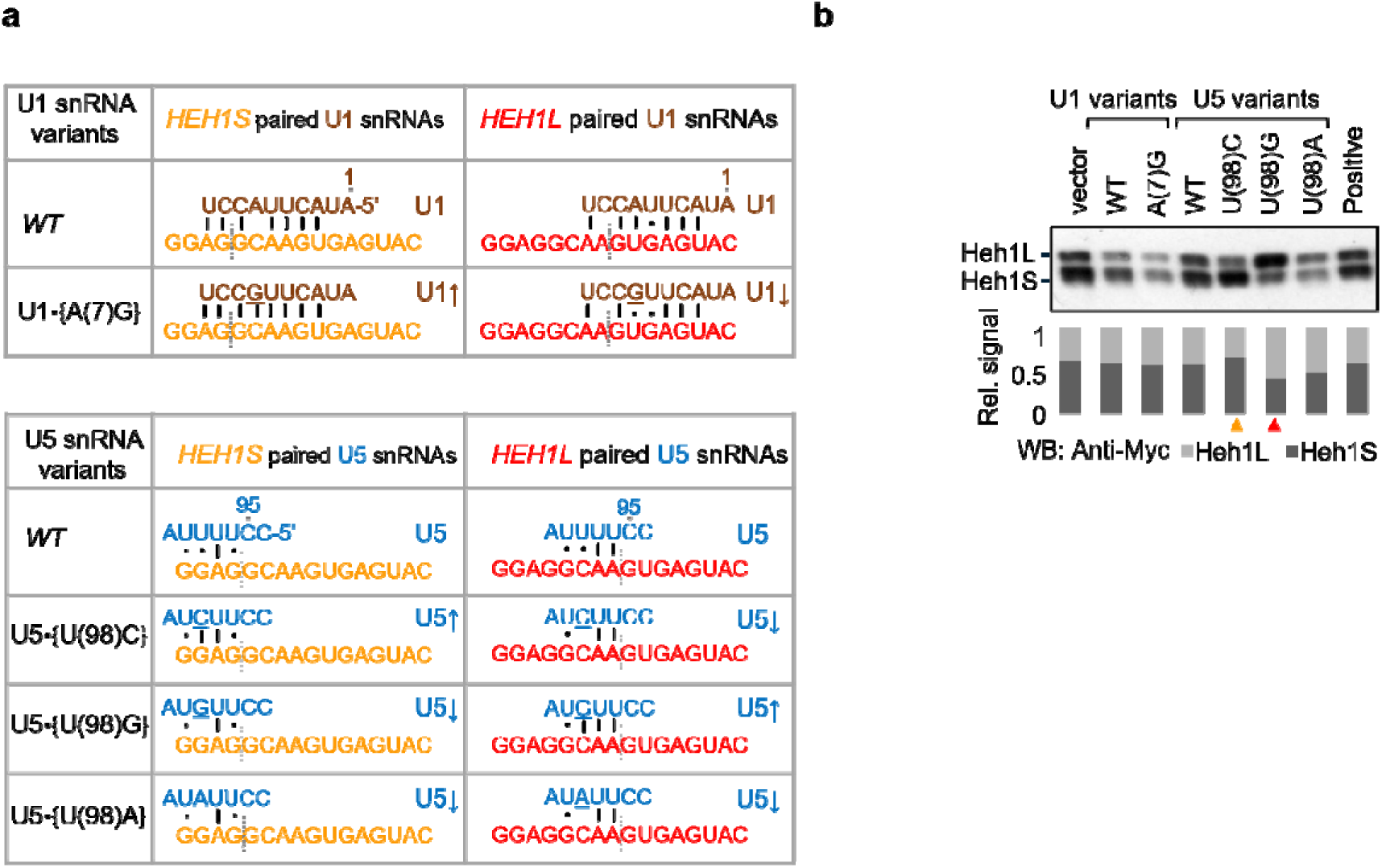
Overexpression of snRNAs and their variants. **(a-b)** Overexpression of U1 and U5 snRNAs. **(a)** To check the effect of trans-acting snRNAs on alternative 5’SS selection, *WT* U1 snRNA, U5 snRNA, and their variants were cloned in episomal (2μ) plasmids and were compared based on their pairing to *HEH1* alternative sites. The variants were made to stabilize the pairing of U1 snRNA to the 5’ exon/intron sequence of *HEH1* pre-mRNA and U5 loop1 to the 5’ exon. **(b)** The influence of overexpressing snRNA variants on HEH1 alternative splicing was monitored using western blots. Orange and red colored upward triangles (▲) indicate increased Heh1S and Heh1L proteins upon overexpression of U5 variants. Overexpression of U1 snRNA did not show visible changes in alternative 5’SS selection.

**Figure S3:**
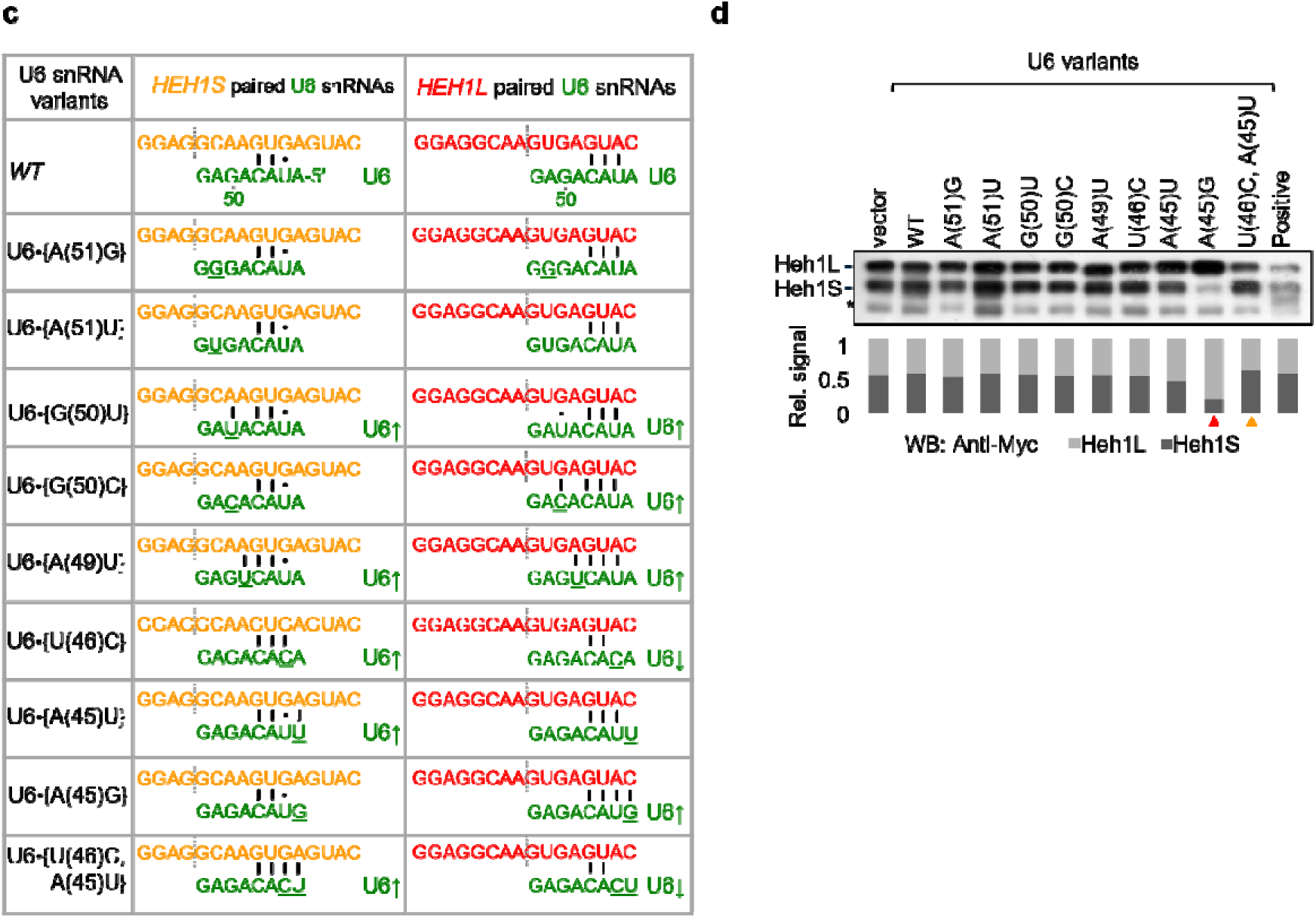
Overexpression of snRNAs and their variants. **(c-d)** Overexpression of U6 snRNA. **(c)** U6 snRNA and its variants are expressed from episomal (2μ) plasmids. The changes were introduced to alter their pairing to the alternative 5’SS. All these variants were made based on extended pairings of U6 ACAGAGAUA to the 5’SS. **(d)** Their influence on *HEH1* alternative splicing was monitored by Western blots. Orange and red upward triangles (▴) indicate variants with higher levels of Heh1S and Heh1L proteins, respectively.

**Figure S4:**
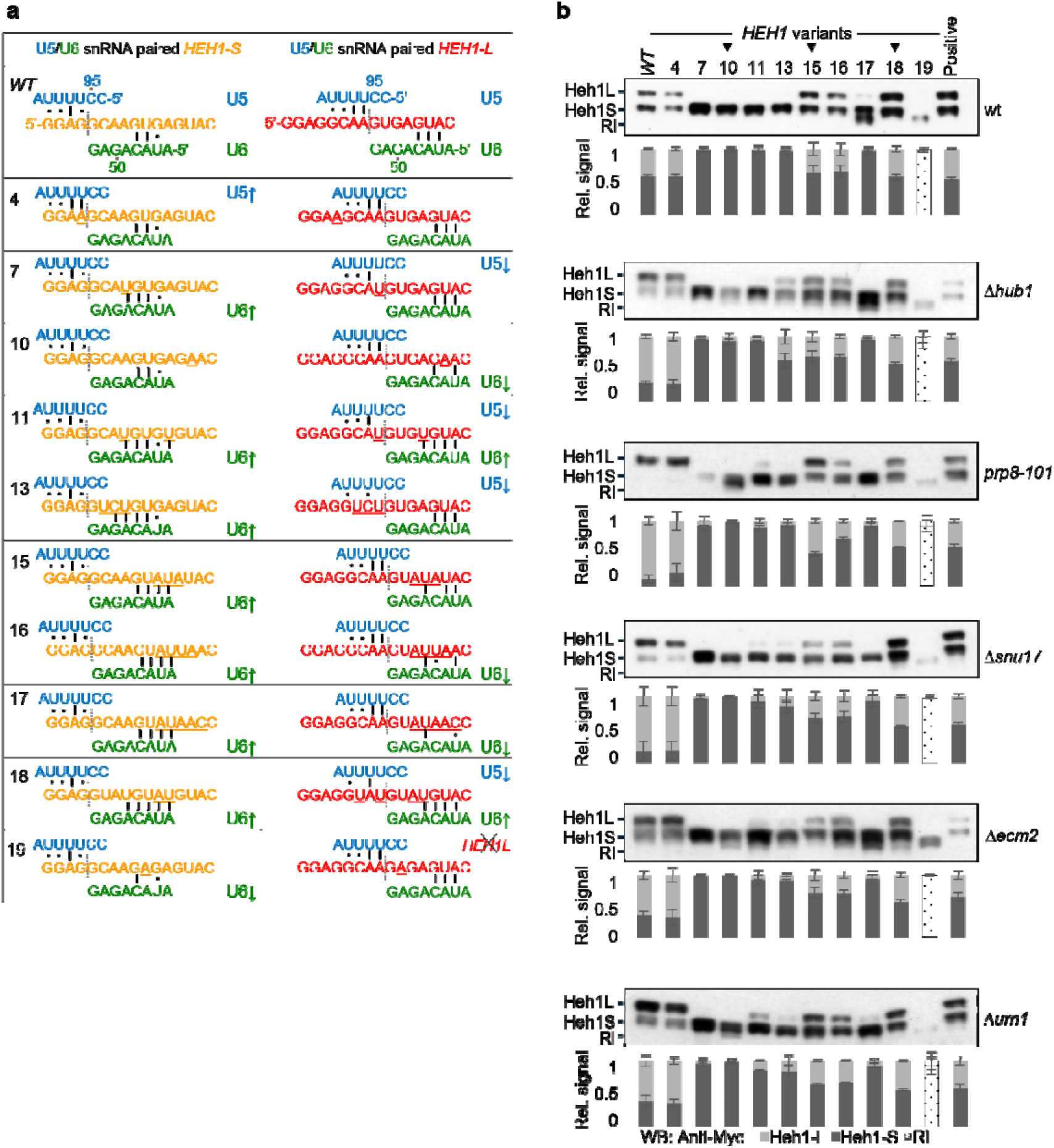
Alternative splicing of *HEH1* pre-mRNA variants. **(a-b)** Assay in the protein factors mutant. **(a)** *HEH1* variants with functional *HEH1* 5’SS splicing was investigated. The binding of U5 and U6 snRNA was compared between the WT and selected variants. x indicates that *HEH1L* 5’SS is not functional in the variant. **(b)** Splicing efficiency of selected variants was monitored by Western blot. Data shown as mean ± s.d. (n=3 independent repeats). RI represents the protein band translated from *HEH1* pre-mRNA with a retained intron.

**Figure S4:**
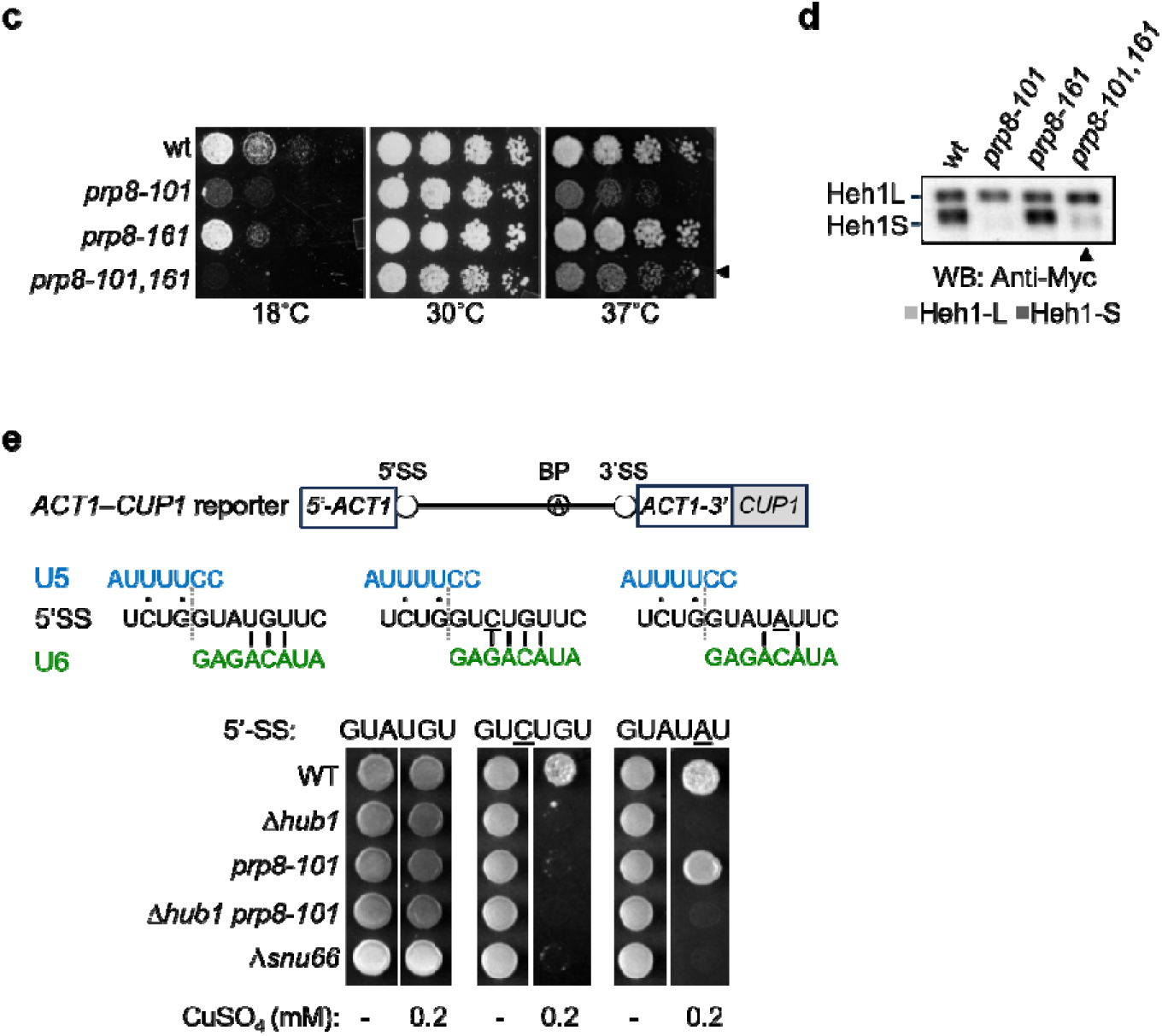
Alternative splicing defect in *prp8-101*. **(c-e)** Combining an opposing allele of *prp8* does not rescue the alternative splicing defect in *prp8-101*. **(c)** The Prp8 alleles, first step allele *prp8-101*(E1960K), second step allele, *prp8-161*(P986T), and both first and second alleles combined, *prp8-101,161,* were spotted and their growth was monitored at 18°C, 30°C, and 37°C. The left triangle (◄) marks the lack of rescue of the growth defect on combining the opposing allele. **(d)** The alleles were tested for *HEH1* alternative splicing by western blot. Combining the first step and the second step allele did not rescue the *HEH1* alternative splicing defect. The up triangle (▲) marks the lack of Heh1S in *prp8-101,161* **(e).** The *ACT1*–*CUP1*^4^ reporter was used to monitor splicing efficiency from non-canonical 5’SS /GUCUGU (also called A3C) and /GUAUAU (also called G5U) in *Δhub1*, *Δsnu66,* and *prp8-101* by growth at different CuSO_4_ concentrations. The mutants showed defective splicing from both non-canonical 5’SS. *prp8-101* showed defects for A3C but not G5U.

**Figure S4:**
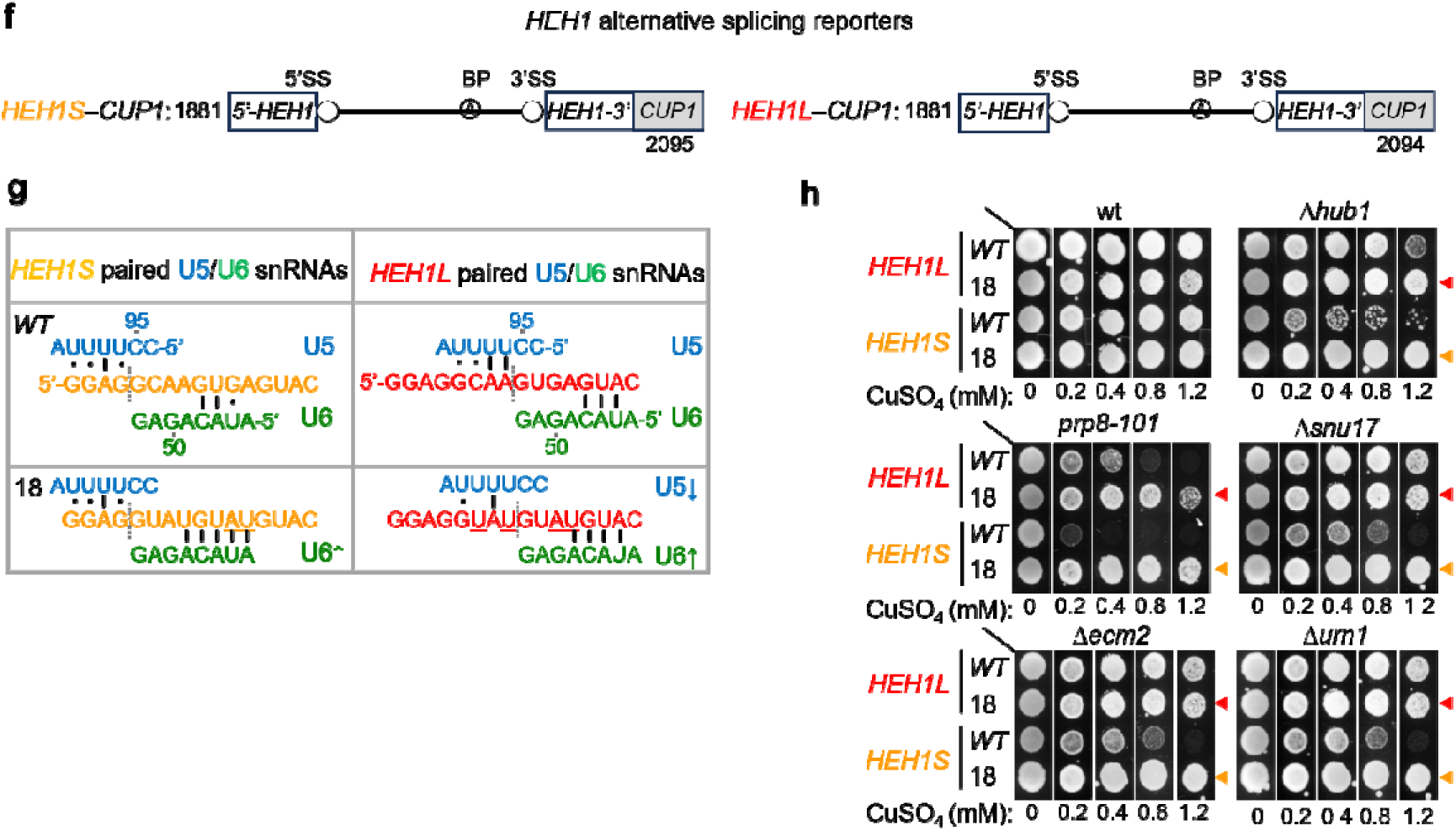
Alternative splicing from the canonical 5’SS does not need the protein factors. **(f-h)** The factors are redundant for alternative splicing from the canonical 5’SS. **(f)** Schematic representation of *HEH1S*–*CUP1 and HEH1L*–*CUP1* reporter. **(g)** Variant 18 /GUAU/GUAUGU was made for both *HEH1S*–*CUP1 and HEH1L*–*CUP1* reporters, and its U5 and U6 pairing was compared**. (h)** The reporter variants were tested for their splicing efficiency at indicated CuSO_4_ concentrations (in mM). The orange and red colored left-pointed triangles (◄) indicate the fully restored growth indicative of normal splicing for *HEH1S*–*CUP1* and *HEH1L*–*CUP1* variants with canonical 5’SS. The reduced growth with *HEH1* 5’SS shows that suboptimal/non-canonical 5’SS have been chosen for its alternative splicing.

**Figure S5:**
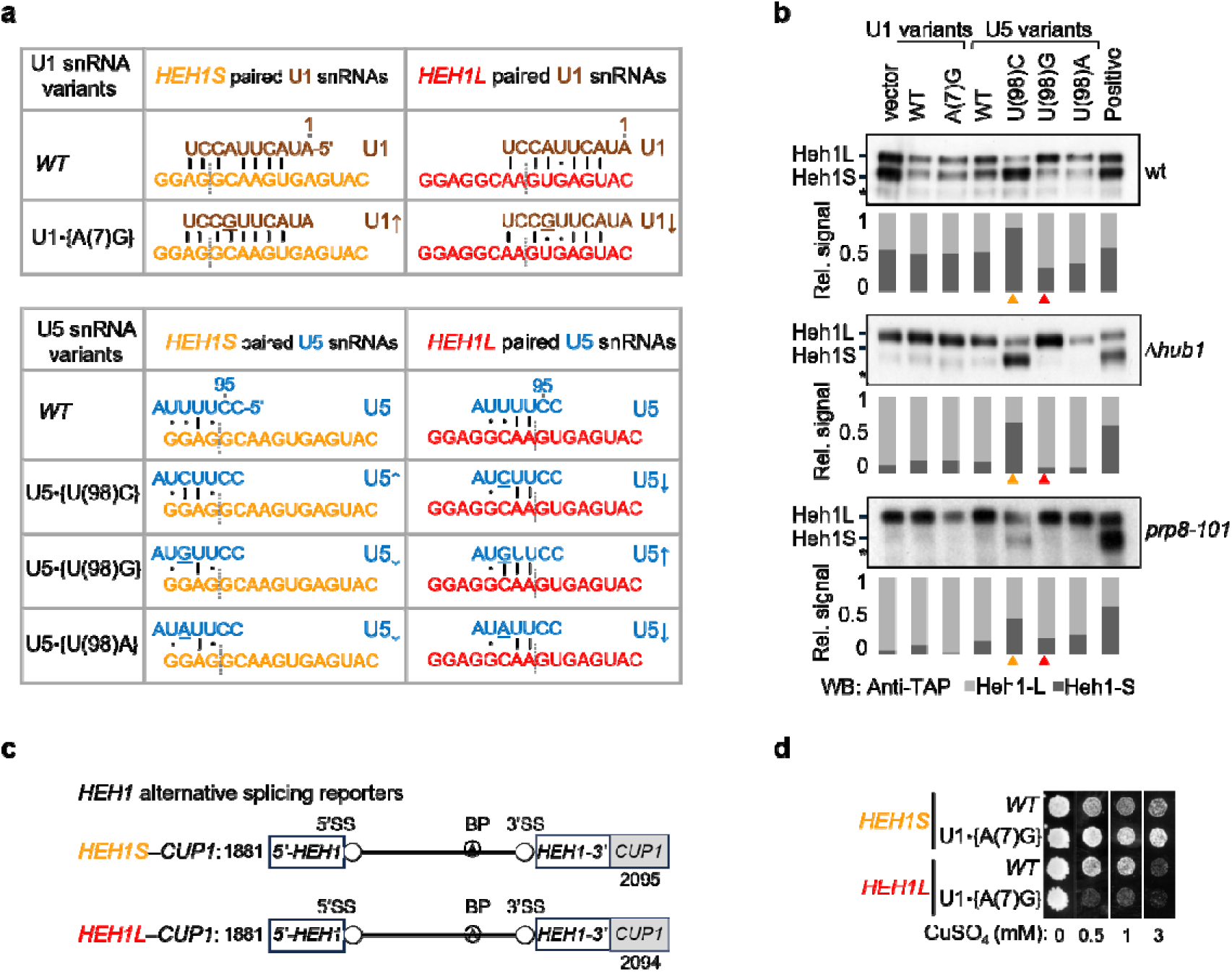
*HEH1* alternative splicing in Δ*hub1 and prp8-101*. **(a-d)** Upon over-expression of U1 and U5 snRNAs variants. **(a)** The pairing of U1 and U5 snRNA variants to *HEH1* alternative isoforms. Upward/downward arrows (↑/↓) indicate an increase/decrease in U5 / U6 pairing compared to *WT* snRNA. **(b)** All the variants were tested for their effect on splicing in wt, *Δhub1,* and *prp8-101* by western blot. Orange and red color upward triangles (▴) indicate higher levels of Heh1S and Heh1L, respectively. **(c)** Schematics of *HEH1-CUP1* alternative splicing reporters. U5 snRNA U(98)C variant enhances selection of *HEH1S* 5’SS, and U(98)G variant enhances selection of *HEH1L* 5’SS. **(d)** *HEH1* alternative splicing reporters and U1snRNA variant, U1 snRNA *WT*, and A(7)G variant. were co-transformed in the wt strain and spotted on indicated CuSO_4_ concentrations. U1 snRNA A(7)G enhances the U1 pairing to *HEH1S* 5’SS and reduces it to *HEH1L* 5’SS. The enhanced growth in *HEH1S*–*CUP1* and poor growth in *HEH1L*–*CUP1* upon U1•{A(7)G expression indicate that this variant showed expected defects in *HEH1L* 5’SS selection, but could not enhance selection of *HEH1S* 5’SS.

**Figure S5:**
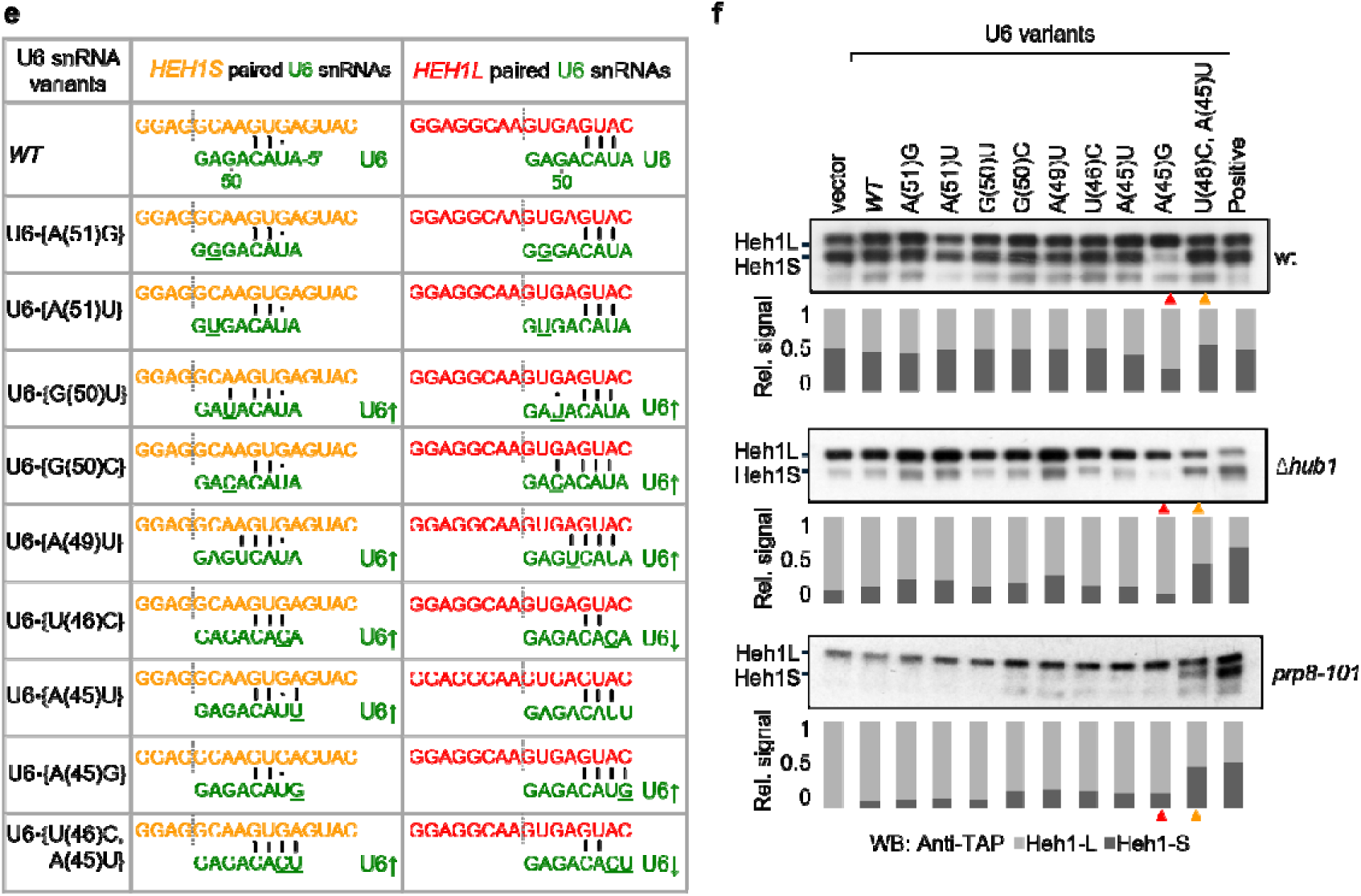
*HEH1* alternative splicing in Δ*hub1 and prp8-101*. **(e-f)** Upon over-expression of U6 snRNAs variants. **(e)** Pairing of U6 snRNA variants to *HEH1S* and *HEH1L* 5’SS. Upward/downward arrows indicate (↑/↓) increase/decrease in U6 snRNA pairing strength. **(f)** All the variants were tested for their effect on splicing in wt, *Δhub1,* and *prp8-101* by western blot. Orange and red upward triangles (▲) indicate a higher amount of Heh1S and Heh1L, respectively. A(45)G variant enhances selection of *HEH1L* 5’SS and U(46)C,A(45)U variant enhances selection of *HEH1S* 5’SS.

